# RBP4 functions as an antagonistic ligand of orphan DR6

**DOI:** 10.1101/2024.10.21.619555

**Authors:** Taeyun Kim, Wonhee John Lee, Thy N. C. Nguyen, Juhee Jang, Kiwook Kim, Min-Sik Kim, Daeha Seo, Young-Sam Lee, Yun-Il Lee

**Affiliations:** Department of New Biology, Daegu Gyeongbuk Institute of Science and Technology (DGIST), Daegu 42988, Republic of Korea; Well Aging Research Center, DGIST, Daegu 42988, Republic of Korea; Department of Physics and Chemistry, DGIST, Daegu 42988, Republic of Korea; Division of Biotechnology, DGIST, Daegu 42988, Republic of Korea

**Keywords:** Death receptor, DR6, orphan receptor, RBP4, Cell death, Calcium ion, Proximity labeling

## Abstract

Death receptor (DR) is a unique transmembrane receptor mediating extrinsic factor-induced programmed cell death pathways. For most DRs, the regulatory mechanism has been revealed to involve agonistic death ligand (e.g., TRAIL, TNF-α)-driven receptor clustering and activation, offering an in-depth understanding of the mechanistic basis for cell death regulation and therapeutic strategies. However, only death receptor 6 (DR6) remains orphan, and its molecular mechanism is poorly understood. Here, we identified retinol binding protein 4 (RBP4), a major vitamin-A carrier protein, as an antagonistic ligand of DR6. Single-molecule photobleaching and single-receptor tracking analyses revealed that RBP4 drives DR6 dimerization on the plasma membrane to prevent DR6 prodegenerative heterointeractions and consequent neurodegenerative processes. DR6 homodimerization predominantly depends on an extracellular intrinsically disordered region, allowing the agonistic propensity of increasing Ca^2+^ levels. Collectively, our study establishes the distinct molecular mechanism of DR6 and highlights the non-canonical role of RBP4 as a cell death regulator.

## Introduction

The extrinsic regulation of the cell death process is predominantly mediated by death receptor (DR) signaling. DRs comprise a subset of the tumor necrosis factor receptor (TNFR) superfamily (TNFRSF) characterized by their cytoplasmic death domains.^1,2^ DR signaling plays pivotal roles in tissue homeostasis and immune function, and its dysregulation contributes to various disease pathologies, including autoimmune diseases, cancer progression and metastasis, and neurodegenerative disorders.^3–7^ The molecular mechanism of DR signaling activation has been understood as a unified model in which the agonistic death ligand drives DR oligomerization and the concomitant formation of the cytoplasmic Fas-associated death domain or TNFR1-associated death domain protein complex, which directly activates downstream cell death pathways.^8^ The findings on these ligands and molecular mechanisms are crucial for comprehending the biological processes depending on the programmed cell death, and more recently, have been used to develop therapeutic strategies to manage the aberrant inflammatory responses and cell loss or to induce targeted elimination of cancer cells.^9–12^

For several decades, the ligand and molecular mechanism of signaling activation have been extensively identified for most DRs. However, only DR6, also known as *Tnfrsf21*, remains an orphan among DRs and TNFRSFs (27 members including DRs), and its mechanism is incompletely understood.^13^ DR6 is globally expressed, and it plays crucial roles in various biological processes, such as the negative regulation of T and B cell function^14–16^, tumor cell proliferation and metastasis^17,18^, and necroptotic and pyroptotic cell death.^18,19^ In particular, functional studies on DR6 mainly focused on the nervous system in which DR6 is highly expressed in mature neurons and oligodendrocyte lineages. Recent studies suggested that DR6 is a critical determinant of neuronal self-destruction during development, injury response, and synaptic plasticity.^20–26^ DR6 also negatively influences oligodendrocyte survival and Schwann cell proliferation, affecting central and peripheral myelination.^27,28^ As DR6 expression is upregulated in patients with Alzheimer’s disease (AD)^29^, amyotrophic lateral sclerosis (ALS)^30^, and Down syndrome^31^, blocking its signaling results in neuroprotective effects against amyloid-beta (Aβ)-induced neurotoxicity and in rodent models of ALS and multiple sclerosis (MS).^27,29,30,32^ This suggests DR6’s central role in neurodegenerative consequences. However, despite continuous findings on the role of DR6 in various cell death processes, the limited knowledge of its signaling regulation largely restricts our understanding of the biological context that governs the prodegenerative process and its further translation into therapeutic applications.

Considering the multiple ligand-binding ability and diversified responses depending on the cellular context of receptor signaling, ligand-dependent receptor signaling is believed to have a complex nature, constructing intricate signaling regulatory networks.^33^ However, studies on the regulatory mechanism of receptors remain challenging because of their transient interaction and complicated factors affecting their dynamics, such as lipid composition and combinatorial cofactors, which consequently leads to the high false-positive rate of traditional methods (e.g., co-immunoprecipitation) and even recent approaches (e.g., high-throughput screening).^34–36^ Thus, a methodology to unwind the complexity of unexplored ligand-receptor networks remains an unmet need.

Proximity labeling uses an enzyme (BioID or APEX) that generates reactive and unstable biotin derivatives and covalently biotinylates biomolecules under close proximity (10–20 nm).^37^ Proximity labeling has been considered to have limited spatial delimitation owing to its rapid and relatively uncontrolled reaction.^37,38^ Thus, it has been mainly utilized for spatial proteomic profiling within compartmentalized organelles (e.g., lysosomes, mitochondria). Its applicability for ligand search in open and crowded extracellular compartments remains unclear.^39^

Here, we show that extracellular proximity labeling (ePL) through N-terminal linkage of TurboID to DR6 specifically enriches endogenous ligand candidates and unveil RBP4 as a ligand of DR6. RBP4 functions as an antagonistic ligand of DR6 by preventing cell death sensitization induced by DR6 overexpression and by blocking the heterotypic prodegenerative interactions of DR6 with amyloid precursor protein (APP) and neurotrophin receptor p75 (p75^NTR^). Through single-molecule analyses, we confirmed that RBP4-induced DR6 dimerization is a crucial event for the antagonistic action of RBP4. The DR6 extracellular domain (ECD) features an exceptionally long intrinsically disordered region (IDR), and we found that native and RBP4-induced DR6 dimerization predominantly depends on the short IDRs adjacent to the RBP4 binding site. Based on these structural properties, we further found that increased extracellular Ca^2+^ content, a hallmark of neuronal stress and neurodegenerative disorders, is the agonistic stimulus interfering with DR6 oligomerization and promoting prodegenerative interactions. Last, we confirmed the functionality of the RBP4–DR6 axis via the protective effect of RBP4 in nerve growth factor (NGF)-deprived sensory axon pruning and Aβ-induced cortical neuronal death models, both characterized as DR6 prodegenerative interaction-dependent. Consequently, we deorphanized DR6, the last orphan death receptor, and established the non-canonical death receptor mechanism in which the antagonistic ligand controls the salt-induced agonistic propensity.

## Results

### ePL specifically enriches endogenous DR6 ligand candidates

We designed a straightforward system in which TurboID, an engineered version of BioID^40^, is linked to the N-terminus of DR6 via a flexible (GGGGS)_2_ linker (Figure 1A). The conditioned media could contain cis or trans regulatory proteins (Figure 1A: left and right, respectively). For comparison, we generated negative controls in which ECD was substituted with enhanced green fluorescent protein (EGFP) or deleted. DR6 is known to undergo heavy glycosylation (N– and O-linked), resulting in multiple shifted bands on western blotting (WB).^41^ TurboID-DR6 also exhibited multiple shifted bands from its native size, indicating conserved glycosylation (Figure S1A). Compared with other DR members, ectopic expression of DR6 conditionally led to cell death in a context-dependent manner.^41^ Similarly, ectopic expression of DR6 or TurboID-DR6 did not induce cell death but sensitized cells to death triggered through mild treatment with sodium arsenite, a toxin that causes mitochondrial oxidative stress and motor neuron death in a DR6-dependent manner (Figure S1B and S1C).^30^ This effect was abrogated by deletion of the cytoplasmic death domain, suggesting engagement of death domain signaling activation (Figure S1C). Therefore, we confirmed that TurboID conserved DR6 functionality.

**Fig. 1.**
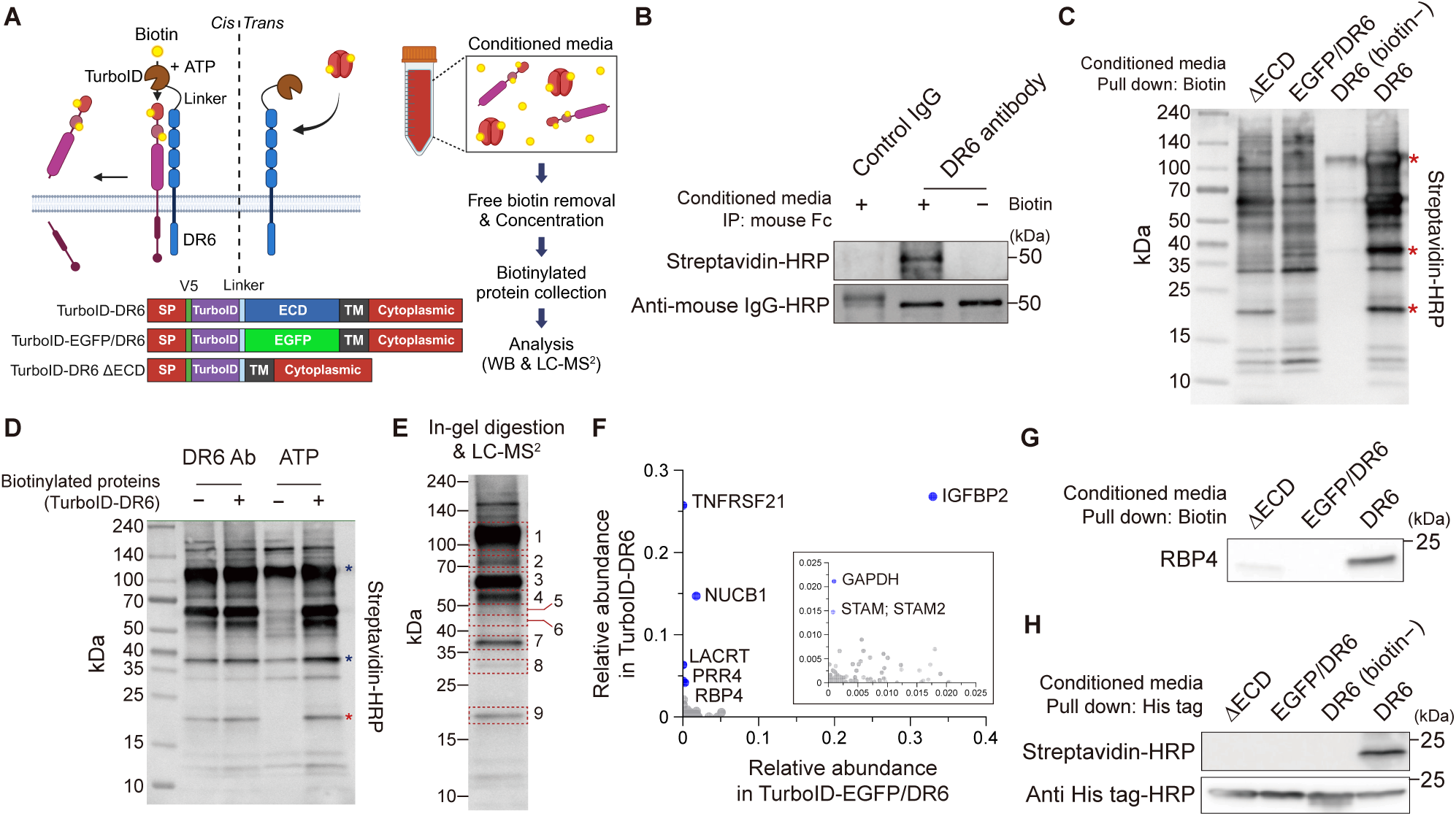
Scheme for DR6 ligand search system utilizing proximity labeling. (**A**) Scheme depicting the DR6 ligand search system utilizing N-terminal TurboID linkage. The biotinylated proteins in conditioned media may contain the proteolytic fragment of *cis*-interacting proteins or *trans* secretory molecules. (**B**) WB showing the specific biotinylation of DR6 antibody compared to control mouse IgG by TurboID-DR6. (**C**) WB of biotinylated proteins collected from the conditioned media from SH-SY5Y transfected with TurboID-DR6 or negative controls. DR6-specific proteins were indicated as a red asterisk. (**D**) WB of biotinylated proteins collected from TurboID-DR6 expressing SH-SY5Y in the presence of DR6 antibody (lane 1, 2) or absence of ATP (lane 3, 4) in the media. A red asterisk indicates a highly ATP-dependent protein. A yellow asterisk indicates partial dependency on ATP. (**E**) The size information of gel cuts subjected to LC-MS^2^. Size-matched gels from TurboID-EGFP/DR6 were used for comparison. (**F**) The relative iBAQ level of each protein in TurboID-EGFP/DR6 and TurboID-DR6. TNFRSF21 (DR6), LACRT, and PRR4 were highly and exclusively enriched in TurboID-DR6 sample. (**G**) Validation of specific enrichment of RBP4 in the biotinylated proteins set collected by TurboID-DR6. (**H**) Confirmation of direct biotinylation of RBP4-His by TurboID-DR6.

No cytotoxic effects were observed with extrinsic supplementation of 1 mM adenosine triphosphate (ATP) and stable expression of TurboID-DR6. Assessing TurboID-DR6’s potential biotinylation of extrinsic binding partners revealed that it biotinylated extracellular DR6 antibodies with high specificity against control mouse immunoglobulin G (IgG) (Figure 1B and Figure S1D), demonstrating that binding proteins can be specifically biotinylated, even in the accessible yet densely populated extracellular environment.

Through this system, we identified several proteins specifically enriched in a DR6 ECD-dependent manner (Figure 1C). Although sodium arsenite markedly altered the biotinylation profile, it mainly inhibited rather than generated biotinylation of new proteins (Figure S1E). Moreover, biotinylation in Hs683, an oligodendroglioma cell line, was predominantly observed in larger proteins, highlighting the occurrence of cell type-specific protein enrichment (Figure S1F). To determine whether these proteins were biotinylated in the extracellular compartment, we used DR6 antibody treatment or omitted ATP from the media (Figure 1D). Biotinylation remained unchanged following DR6 antibody treatment but was ATP presence-dependent. ATP serves as a membrane-impermeable source for generating biotinyl-5-AMP via TurboID. Thus, these findings indicate that DR6-specific proteins were biotinylated in the extracellular environment, establishing them as DR6 ligand candidates.

### RBP4 is a ligand candidate of DR6

To identify candidate proteins, in-gel digested proteins were analyzed using mass spectrometry. Protein-enriched and size-matched gel cuts (nine each) from TurboID-EGFP/DR6 and TurboID-DR6 samples were subjected to liquid chromatography with tandem mass spectrometry (LC-MS^2^) (Figure 1E). Consequently, 120 secretory proteins were identified of which 50 secretory proteins were enriched in the TurboID-DR6 sample (Figure S2A and S2B; Supplementary Table 1). Several proteins, such as proline-rich protein 4 (PRR4) and lacritin (LACRT), were specifically enriched in TurboID-DR6 samples, whereas insulin-like growth factor-binding protein 2 (IGFBP2) was the most abundant protein in TurboID-EGFP/DR6 and TurboID-DR6 samples (Figure 1F). The relatively low proportion of TurboID-DR6–specific proteins might be attributable to the clustering of TurboID-EGFP/DR6 with endogenous DR6 via transmembrane domain interactions or subcellular localization guided by signal peptides. Similarly, highly abundant proteins in TurboID-DR6 samples, such as RBP4, glyceraldehyde 3-phosphate dehydrogenase (GAPDH), and signal-transducing adapter molecule (STAM), were also rarely found in TurboID-EGFP/DR6 samples (Figure 1F). Considering such innate non-specificity, we analyzed all TurboID-DR6– enriched secretory proteins to specify ligand candidates.

The secretome featured by TurboID-DR6 projects the molecular landscape of amyloidosis and AD (Figure S2C–S2E). DR6 is known to interact with APP in cis to execute axon pruning by trophic factor (TF) deprivation^42,43^ and we found specific enrichment of APP by TurboID-DR6 (Figure S2F). Network analysis of proteins enriched by TurboID-DR6 revealed an APP-centered interaction network (Figure S2G). However, additional AD-associated signaling proteins (IGFBP2, transforming growth factor β1 (TGFB1), and RBP4) were also enriched, suggesting a probable APP-independent link between DR6 and AD-associated signaling molecules.

Recent studies revealed significant DR6 upregulation in the brains of patients with AD, and DR6 signaling mediates TF deprivation-induced axon pruning and Aβ-induced neurotoxicity, which are cognate models of AD pathology. Thus, to identify a functional ligand that links DR6 signaling to AD-associated neuropathies, we assessed the enrichment of AD-associated secretory proteins via WB (Figures 1G and S2H) and revealed that RBP4 was specifically collected and directly biotinylated by TurboID-DR6 (Figure 1H).

RBP4, a major carrier of retinol, is primarily expressed in the liver, the main organ for retinol storage. Transthyretin (TTR) specifically clusters with retinol-bound RBP4 (holo-RBP4), not with apo-RBP4, facilitating RBP4 secretion from hepatocytes and preventing renal clearance; thus, holo-RBP4 is dominant in circulation.^44^ Although RBP4 has been primarily studied in metabolic contexts and disorders owing to its major role in retinol delivery, a growing body of evidence indicates its close association with diverse neurodegenerative disorders. Notably, RBP4 can cross the blood–brain barrier (BBB), and serum RBP4 levels have been inversely correlated with various neuropathies, such as autistic regression^45^, ALS prognosis rate^46^, diabetic mild cognitive impairment^47^, and brain volume loss in patients with MS^48^. Additionally, RBP4 levels in cerebrospinal fluid, but not in plasma, have been shown to gradually decrease as AD progresses^49–51^. Furthermore, genetic depletion of RBP4 results in a broad range of neuropathies, including impaired y-maze performance, compromised locomotor activity, and neocortical and hippocampal neuronal loss, mechanically implicating reduced RBP4 levels in neurodegeneration.^52^ However, despite its crucial relationship with neurodegenerative disorders, the detailed mechanistic link between RBP4 depletion and neuropathies remains unknown.

### RBP4 directly interacts with the DR6 ECD

The direct interaction between recombinant DR6 ECD and RBP4 was confirmed via surface plasmon resonance (SPR) analysis (Figure 2A). The interaction profile exhibited an almost “stable” pattern characterized by slow dissociation. To further characterize the DR6–RBP4 interaction, we crosslinked recombinant RBP4 and the DR6 ECD using bis(sulfosuccinimidyl)suberate (BS^3^). The DR6–RBP4 crosslinked complex predominantly occurred with 1:1 stoichiometry, as indicated via WB (Figure S3A). Subsequently, the crosslinked sites of in-gel digested proteins were analyzed using LC-MS^2^ (Figure S3B and Supplementary Table 2). A docking simulation, assuming one-to-one binding within the restraints, yielded a single binding model (Figure 2B and Figure S3C). The positively charged surface of DR6, enriched with lysine and arginine, was predicted to interact with RBP4’s polar residues, with hydrophobic interfaces playing a minor role in the interaction (Figure 2C and Figure S3D–S3F). Alanine substitution of these lysine residues significantly altered the interaction with RBP4, as evidenced through the bioluminescence resonance energy transfer assay between 5-Carboxyltetramethylrhodamine (5-TMR)-labeled RBP4 and NanoLuc luciferase (Nluc)-DR6 (NanoBRET; Figure 2D). These findings elucidate the interaction between RBP4 and the DR6 ECD, which utilizes the positively charged surface, a featured region of DR6 (Figure S3G).

**Fig. 2.**
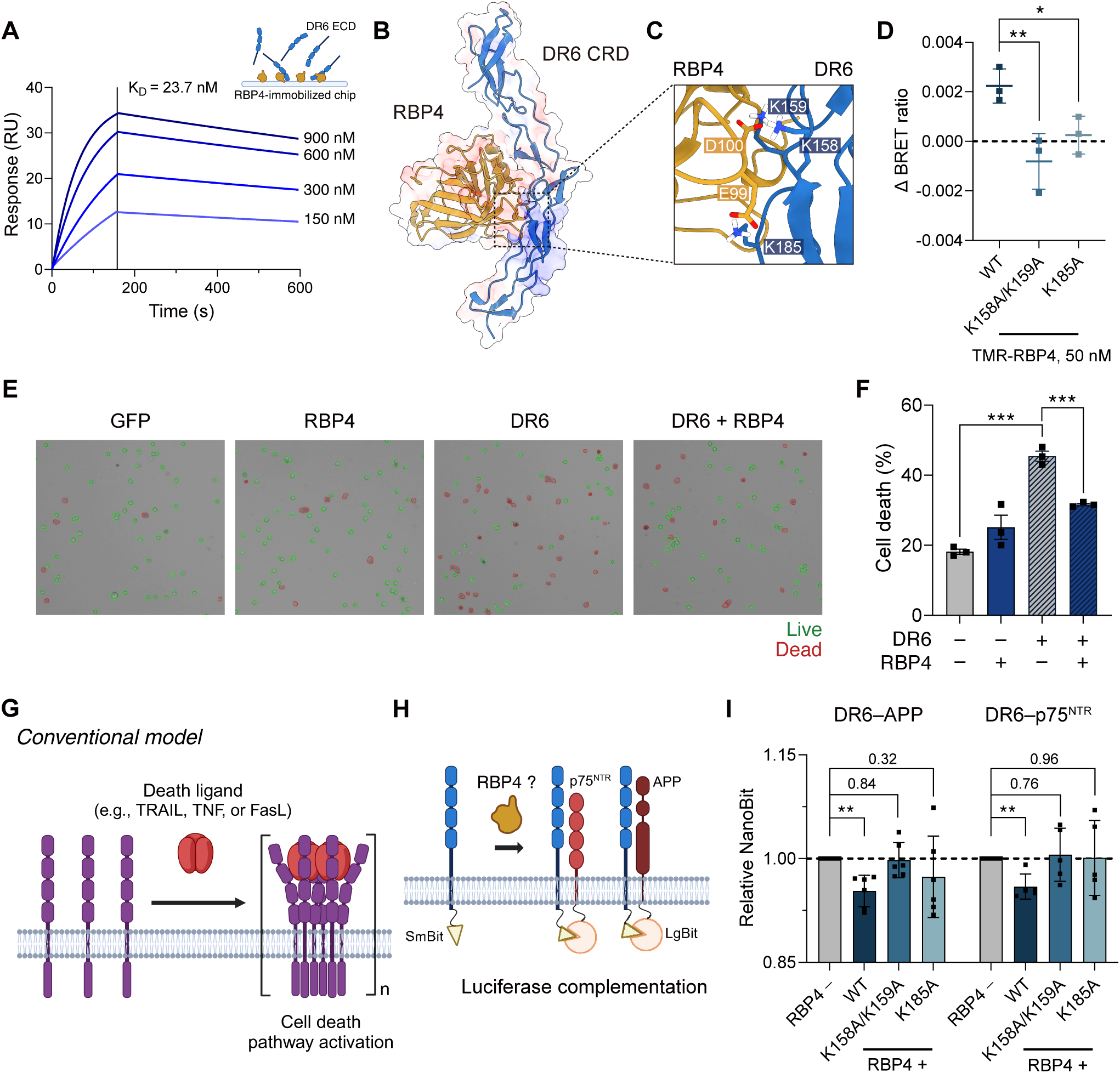
RBP4 directly binds to DR6 and plays an antagonistic role in vitro. (**A**) Surface plasmon resonance analysis on the elevating concentration of recombinant DR6 ECD flew to the RBP4-immobilized sensor chip. (**B**) A possible 1:1 binding structure between RBP4 and DR6 cysteine-rich domain (CRD) simulated with the crosslinked residues as a restraint. A transparently colored surface indicates the charge. (blue=positive; red=negative) (**C**) Close view of the key residues for DR6-RBP4 interaction. (**D**) BRET analysis between 5-TMR-labeled recombinant RBP4 and Nanoluc-linked DR6. Data are presented as mean ± SD determined from three independent experiments performed in quadruplicate. \**p*<0.05 and \*\**p*<0.005 by unpaired two-tailed t-test. (**E**) Representative image showing the automated counting of live/dead cells following the trypan blue staining. (**F**) Quantification of the cell death rate of SH-SY5Y cells transfected with plasmids following the 24-hour treatment of 300 μM sodium arsenite. The cell death rate was automatically counted by a cell counting device. Data are presented as mean ± SD determined from three independent experiments. \*\*\**p*<0.0005 by unpaired two-tailed t-test. (**G**) Scheme depicting conventional agonistic death ligand-dependent DR activation model. (**H**) Scheme depicting NanoBit assay measuring the effect of RBP4 on the heterotypic interactions between DR6 and APP or p75^NTR^. (**I**) NanoBit assay measuring the effect of recombinant RBP4 in DR6 interaction with APP or p75^NTR^. Heterotypic interactions of DR6 wild-type or mutants were measured following the treatment of recombinant RBP4 (0.5 μM) for 30 min. Data are presented as mean ± SD determined from six (APP) or five (p75^NTR^) independent experiments performed in quadruplicate. \*\**p*<0.005 by unpaired two-tailed t-test.

### RBP4 serves as an antagonistic ligand of DR6

We previously observed the enrichment of endogenous RBP4 in the TurboID-DR6 stable cell line without cytotoxicity. Additionally, sodium arsenite inhibited the biotinylation of several candidates through TurboID-DR6 (Figure S1E) but did not induce new protein biotinylation. Notably, we observed that coexpression of RBP4 and DR6 did not induce cell death, but RBP4 significantly impeded sensitization to sodium arsenite-induced cell death in DR6-coexpressing cells (Figure 2E and 2F). Therefore, we hypothesized that RBP4 plays an antagonistic role in DR6 signaling activation.

To date, the identified ligands of DRs have been agonists, with signaling activated through multimeric ligand-driven receptor homotypic clustering and transduced from cytoplasmic death domain oligomerization (Figure 2G). However, evidence for DR6 homo-clustering in signaling activation is lacking, whereas heterotypic interactions with APP or p75^NTR^ are implicated in axon pruning and neuronal cell death. The crystal structure of DR6–APP complex shows that the prodegenerative interaction can occur without the participation of any agonistic molecule. Thus, we hypothesized that DR6 has a distinct molecular mechanism where an antagonistic ligand controls autonomous heterotypic prodegenerative interactions. To test this hypothesis, we examined RBP4’s inhibition of DR6 cis interaction with APP (APP751; NP_958816) and p75^NTR^ through the luciferase complementation assay (NanoBit; Figure 2H). Interestingly, RBP4 significantly inhibited DR6 heterotypic interactions with APP and p75^NTR^, whereas K158A/K159A or K185A mutants abrogated this effect, indicating direct inhibition by RBP4 (Figure 2I). Thus, these findings indicate that RBP4 directly interacts with DR6 ECD and acts as an antagonistic ligand, not a conventional agonistic death ligand.

### RBP4 forms a complex with DR6 in 2:2 stoichiometry

Considering both the simultaneous dual inhibition of the interactions with APP and p75^NTR^ (Figure 2I) and the structural properties of DRs lacking obvious pocket for stable one-to-one ligand interactions and dependency on the multimeric complex formation (Figure 2B and 2G), we hypothesized that RBP4 may forms an “inactive” complex with DR6. Thus, we first sought to determine the stoichiometry of RBP4-DR6 complex.

Owing to the similar and small size of RBP4 and DR6 ECD and the heavy N– and O-linked glycosylation of DR6, typical mass-based protein stoichiometric analyses, such as size exclusion chromatography–multiangle light scattering method, pose challenges (data not shown). Instead, we employed a single-molecule dual-color photobleaching assay, which was recently used for protein stoichiometric analysis (Figure 3A). Consistent with the previous stable interaction profile in SPR (Figure 2A), Alexa Fluor 488-labeled RBP4 and Alexa Fluor 660-labeled DR6 ECD were observed in single-molecular spots and separately bleached in a stepwise manner (Figure 3B and 3C). Determination of the counts of fluorescent dye within the complexed single-molecular spots revealed a selectively increasing dimeric population of DR6 and RBP4 along with colocalization (Figure 3D and 3E). Interestingly, DR6 ECD mainly exhibits a monomeric and minorly populated dimeric state (approximately 10%) (Figure 3D), whereas RBP4 exhibits a native propensity to oligomerize and selectively dimerize during its interaction with DR6 (Figure 3E), indicating RBP4-driven DR6 oligomerization. Within the RBP4–DR6-colocalized populations, monomeric RBP4 was primarily associated with monomeric DR6, whereas dimeric RBP4 was colocalized with dimeric DR6 (Figure 3F). Although infrequently detected, trimeric RBP4 was associated with dimeric DR6 (Figure 3F). Given the selective increase in the dimeric RBP4 population and its association with dimeric DR6, this pattern indicates a dominant population of 2:2 complex. Consistently, the top-ranked model for 2:2 RBP4-DR6 complex generated by Alphafold3 suggests that dimerization is mainly driven by RBP4 (Figure S4A and S4B) and DR6 K159 and K185 play a crucial role in the interaction with RBP4 (Figures 4G and S4C). RBP4 molecules are predicted to rearrange and form a hydrophobic and symmetric RBP4–RBP4 interface (Figure S4D and S4E). Additionally, along with the symmetric RBP4 interface, DR6 hydrophobic residues (proline and tryptophan) are symmetrically aligned, extending a long hydrophobic interface (Figure S4F). This model is consistent with both our findings on RBP4’s specific propensity to oligomerize and previous findings that the DR6 cysteine-rich domains do not dimerize.^53^ Collectively, these data reveal the favorable 2:2 complex formation with implications for the binding structure.

**Fig. 3.**
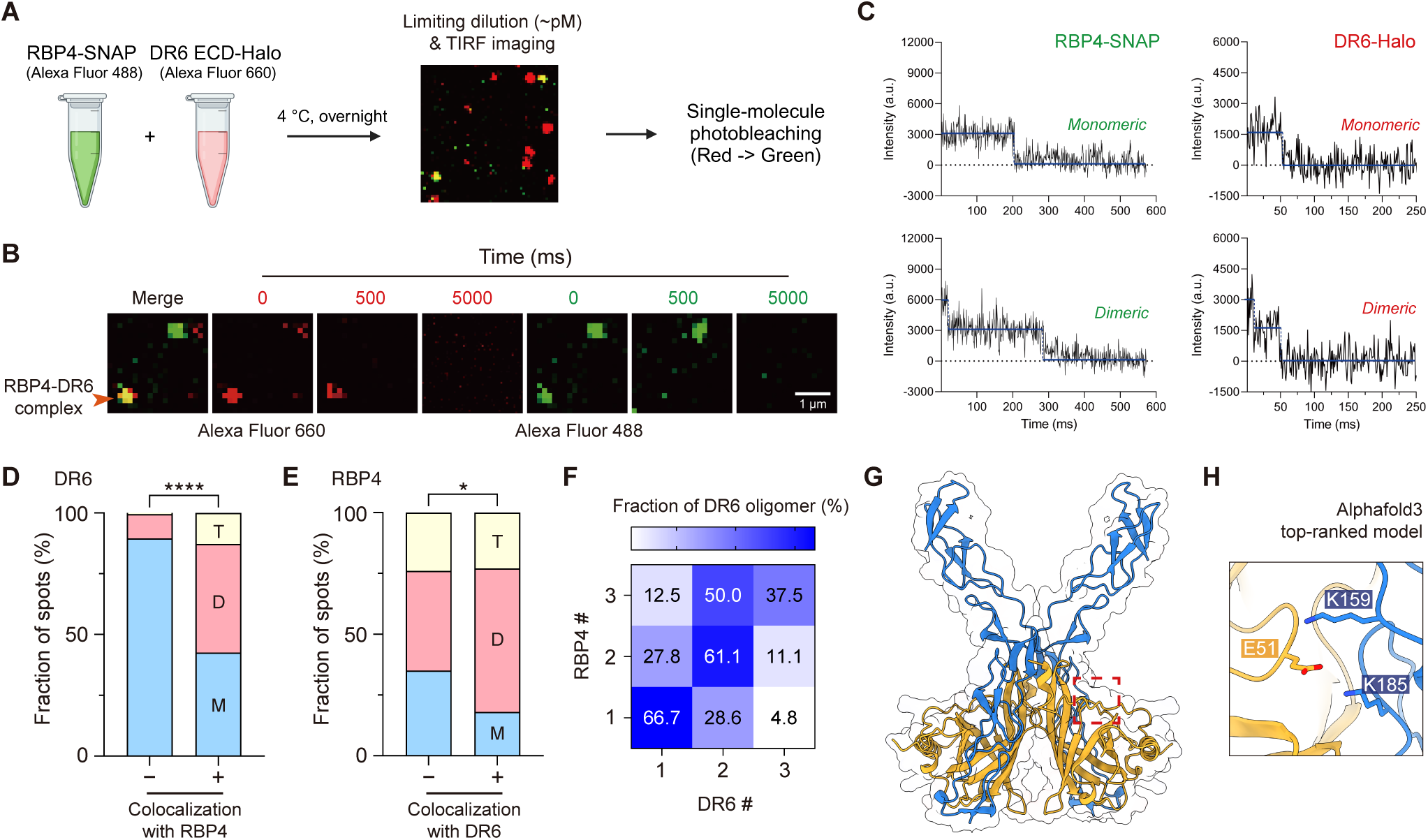
RBP4 forms 2:2 complex with DR6. (**A**) Scheme showing the stoichiometric analysis of RBP4-DR6 complex. (**B**) Representative image showing single-molecular RBP4-SNAP (green) and DR6 ECD-Halo (red). The arrow indicates the colocalized single-molecular spots. (**C**) Representative image showing the stepwise photobleaching profile. (**D** and **E**) Population of DR6 (D; n=211 non-colocalized spots and n=47 colocalized spots) and RBP4 (E; n=210 non-colocalized spots and n=50 colocalized spots) oligomers with or without colocalization. (T=trimer, D=dimer, and M=monomer) \**p*<0.05 and \*\*\*\**p*<0.0001 by unpaired two-tailed chisquared test. (**F**) The ratio of colocalized DR6 oligomers within the RBP4 oligomeric state (mono, di, or trimer). (**G**) Top-ranked structural model for 2:2 complex predicted by Alphafold3. (**H**) Close view of DR6 K159 and K185 predicted to interact with RBP4 E51.

**Fig. 4.**
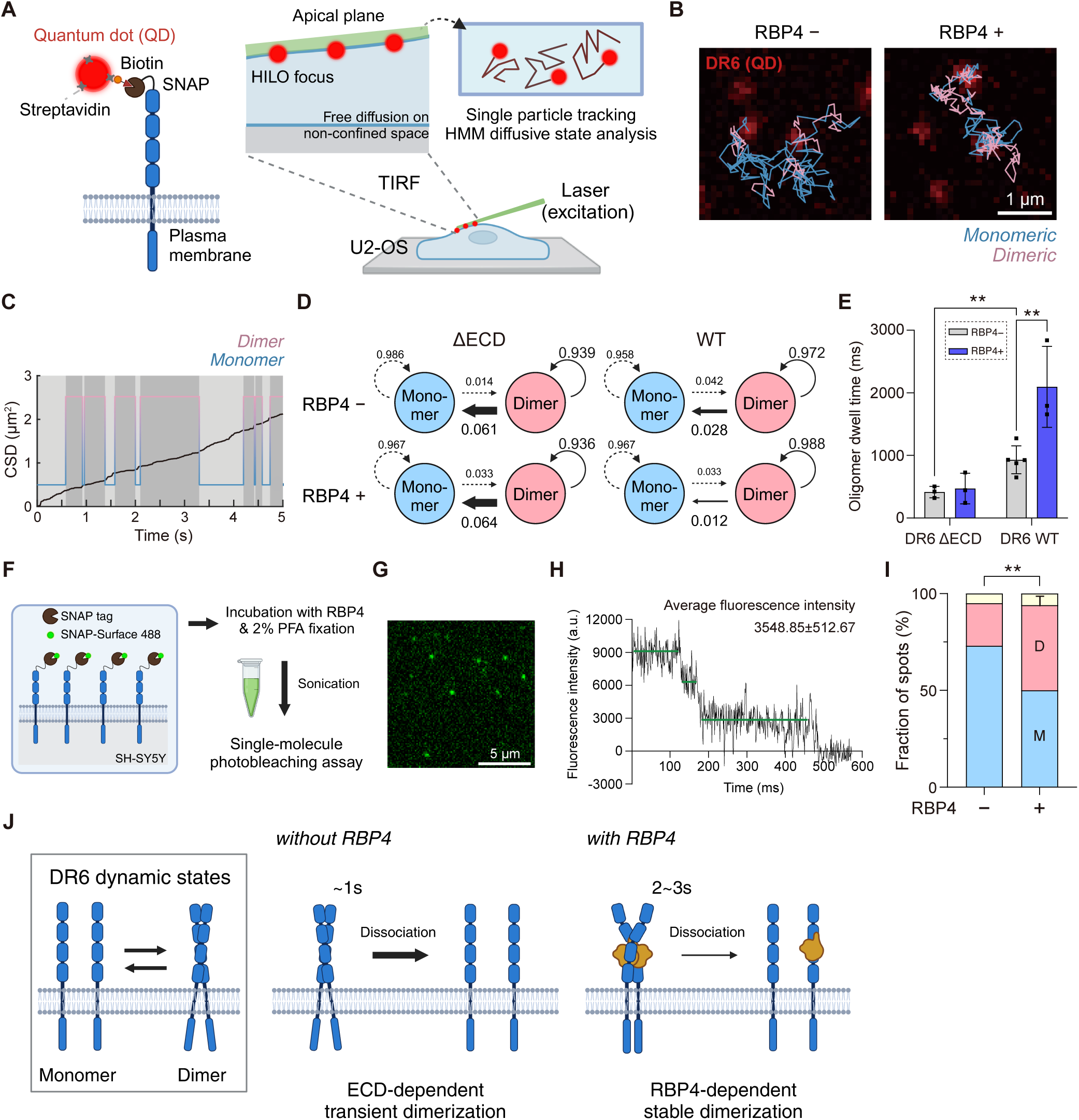
Single-receptor tracking reveals RBP4 induces DR6 dimerization on plasma membrane. (**A**) Scheme depicting QD labeling of DR6 and single-receptor tracking. (**B** and **C**) Representative image showing single DR6 trajectory (B) and oligomeric state analysis (C). (**D**) Hidden Markov chain of diffusive state transition showing the average turnover rate between receptor oligomeric states with or without RBP4. (**E**) The dwell time of oligomeric DR6 wildtype (n=3 cells with RBP4 and n=5 cells without RBP4) and DR6 ΔECD (n=3 cells) with or without RBP4 treatment measured by single-particle tracking of QD-labeled DR6. Data are presented as mean ± SD determined from QD-labeled independent cells. \*\**p*<0.005 by unpaired two-tailed t-test. (**F**) Scheme depicting specific fluorescence labeling and stoichiometric analysis of membrane-bound DR6. (**G**) Representative image showing single-molecular separation of SNAP-DR6. (**H**) Representative image of the stepwise photobleaching profile of SNAP-DR6. (**I**) Population of DR6 with (n=144) or without (n=152) RBP4 exposure. \*\**p*<0.005 by unpaired two-tailed chi-squared test. (**J**) Model for the dissociation of native– and RBP4-bound-DR6 dimer. Time represents the average oligomer dwell time.

### RBP4 drives DR6 dimerization on plasma membrane

Stoichiometric analysis between the recombinant DR6 ECD and RBP4 suggests a model in which RBP4 influences DR6 dimerization. To characterize DR6 dynamics and its alteration by RBP4 on the living cell plasma membrane, we leveraged advancements in single-molecule tracking, enabling quantitative measurement of receptor dynamics and stoichiometry in living cells. Specifically, we characterized DR6 dynamics on the plasma membrane using single-receptor tracking of quantum dot-labeled DR6 (Figure 4A and 4B; Supplementary Video 1– 3). DR6 exhibited two major oligomeric states, with diffusion coefficients close to a monomer (D_m_ = 0.12 μm^2^/s) and a dimer (D_d_ = 0.05 μm^2^/s) (Figure 4C).^54^ Consistent with stoichiometric data, RBP4 significantly decreased the dimer-to-monomer turnover of DR6 (Figure 4D). Monomer-to-dimer transition largely depends on the expression level. Dwell time, representing stability at each state, was calculated based on transitions between oligomeric states. RBP4 significantly increased the dwell time in the dimeric state (Figure 4E). The effect of RBP4 was abrogated in the ECD-deleted DR6 (DR6 ΔECD), indicating that the effect involved an interaction with ECD (Figure 4E). Interestingly, compared with DR6 ΔECD, wild-type (WT) DR6 displayed 2.1-fold lower turnover into the monomeric state and a significantly longer dimeric dwell time, suggesting that the ECD itself drives oligomerization (Figure 4D and 4E). Previous studies on DR5 found that the ECD inhibits receptor clustering that is mainly driven by the interactions between transmembrane domains, indicating different molecular mechanisms for receptor oligomerization between DR5 and DR6.^55^ Thus, through receptor tracking in the living cell plasma membrane, we found that RBP4 promotes DR6 dimerization, which was naturally sustained by the ECD.

To confirm the DR6 homo-dimerization on the plasma membrane, we employed the single-molecule photobleaching assay of SNAP-DR6 expressed in SH-SY5Y labeled with SNAP-Surface Alexa Fluor 488, a membrane-impermeable Alexa Fluor 488 SNAP ligand (Figure 4F). After sonication, the labeled DR6 from crosslinked cell lysates were separately observed in single-molecule level (Figure 4G) and bleached in stepwise manner (Figure 4H). As a result, SNAP-DR6 mainly exhibits a monomeric state, and RBP4 specifically decreased and increased monomeric and dimeric populations, respectively, indicating the dimerization of monomeric DR6 (Figure 4I). Taken together, through a single-molecule analyses, we confirmed that DR6 exhibits relatively transient dimerization inherently sustained by ECD and RBP4 further stabilizes it on the plasma membrane (Figure 4J).

### RBP4-induced DR6 dimerization is crucial for its antagonistic action

Next, we investigated whether RBP4-induced DR6 dimerization is a mechanistic event for the antagonistic action of RBP4. To test this by specifically inhibiting DR6 dimerization without affecting its binding affinity, we first asked the mechanism of the organization of native and RBP4-responsive dimerization, both predominantly driven by ECD. Compared with other DR members, the DR6 ECD features an exceptionally long juxtamembrane IDR, encompassing approximately half of its length (145/308 amino acids in the ECD), predominantly composed of histidine (Figure 5A and S5A–S5C). Using flDPnn^56^, a neural network-based algorithm for computing IDRs and their properties from protein sequences, we consistently identified two short subregions (220–232 and 249–262) with high protein binding propensities (Figure 5B and S5D). These two regions were located adjacent to a cysteine-rich domain (42–215), with amino acid compositions biased toward proline and asparagine/serine, respectively (Figure S5E). PxP (spaced prolines) motif or proline-rich IDRs play a critical role in the formation of biological condensates of proteins and polar residues are hallmarks of prion-like proteins.^57–59^ Therefore, we hypothesized that these short IDRs mediate native and RBP4-responsive DR6 dimerization.

**Fig. 5.**
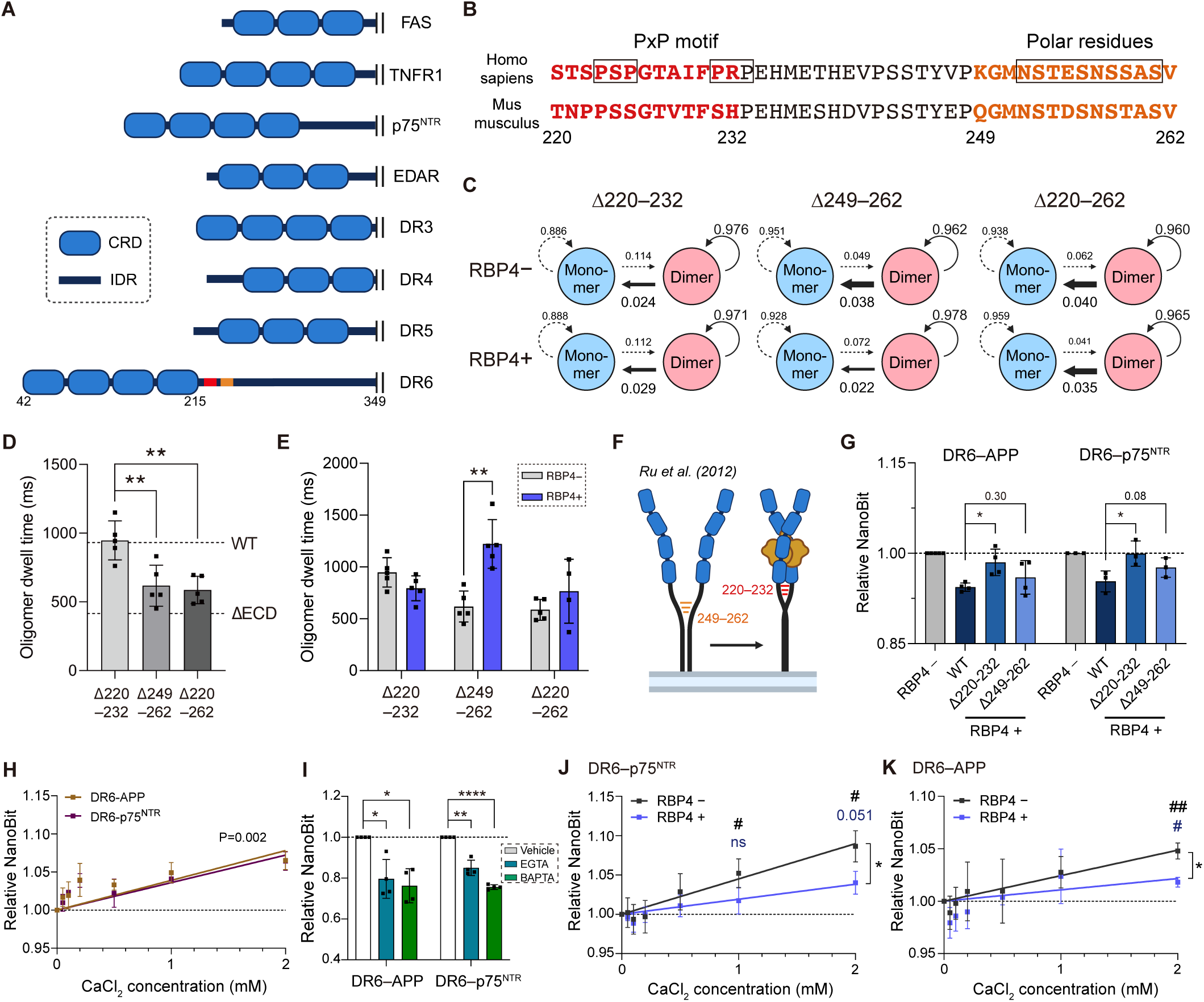
IDR-dependent DR6 dimerization is crucial for the antagonistic action of RBP4 and Ca^2+^ acts as an agonistic stimulus. (**A**) Graphic depicting CRD and IDR in the ECD of DRs. Red and orange regions indicate the subregions (220–232 and 249–262) predicted to have a high protein binding propensity. (**B**) The sequence of subregions. PxP motif and polar residues are indicated in the box. (**C**) Hidden Markov chain of diffusive state transition showing the average turnover rate between receptor oligomeric states of DR6 mutants with or without RBP4. (**D**) Dwell time in the dimeric state of native DR mutants. (n=4) (**E**) Dwell time in the dimeric state of DR6 mutants with or without RBP4 treatment. (n=5 cells except for RBP4-treated Δ220–262 mutant (n=4)) Data of RBP4-non-treated samples are previously presented in (D). Data are presented as mean ± SD determined from QD-labeled independent cells. \*\**p*<0.005 by unpaired two-tailed t-test. (**F**) Graphic depicting the utilization of IDRs in DR6 dimerization. (**G**) NanoBit assay measuring the effect of RBP4 on the interaction of DR6 wildtype or mutants with APP (n=4) or p75^NTR^ (n=3). Data are presented as mean ± SD determined from independent experiments performed in quadruplicate. \**p*<0.05 by unpaired two-tailed t-test (**H** and **I**) NanoBit assay measuring the interaction of DR6 with APP or p75^NTR^ (H) along with the elevating concentration of CaCl_2_ (n=4) and (I) after 10 min incubation with BAPTA (2 mM) or EGTA (5 mM) under 2 mM CaCl_2_ presence (n=4). Data are presented as mean ± SEM determined from four independent experiments performed in quadruplicate. \**p*<0.05, \*\**p*<0.005, and \*\*\*\**p*<0.0001 by one sample t-test. (**J** and **K**) NanoBit assay measuring the effect of the elevating concentration of CaCl_2_ with or without recombinant RBP4 (0.5 μM) in DR6 interaction with p75^NTR^ (J; n=5) and APP (K; n=4). Data are presented as mean ± SEM determined from independent experiments performed in quadruplicate. #*p*<0.05 and ##*p*<0.005 by one sample t-test. \**p*<0.05 by unpaired one-tailed t-test.

Interestingly, using single-receptor tracking, we found that both IDRs play pivotal roles in DR6 dynamics through different mechanisms. IDR deletion did not alter the oligomeric states of DR6 mutants. (Figure 5C). Deletion of asparagine/serine-rich region (249–262) markedly inhibited DR6 native dimerization to a level comparable with DR6 ΔECD, indicating the pivotal role in native dimerization (Figure 5D). In contrast, deletion of the proline-rich region (220–232) completely abolished RBP4-induced dimerization (Figure 5E). These findings indicate that DR6 organizes its dimerization and responsiveness to RBP4 through distinct short IDRs adjacent to the RBP4 binding site (Figure 5F). The dependency on the adjacent IDR is consistent with the results of previous small angle X-ray scattering analyses in which DR6 ECD dimerization extends from and depends on this IDR (Figure 5F).^53^ Thus, we revealed that DR6 utilizes short IDRs for native and RBP4-induced dimerization.

Given the IDR-dependency of DR6 dimerization, we explored the mechanistic link between the DR6 dimerization and signaling regulation governed by RBP4. IDR deletion did not affect DR6’s interactions with APP and p75^NTR^ (data not shown). However, blocking DR6 dimerization through IDR deletion significantly inhibited the effect of RBP4, indicating that RBP4-induced DR6 dimerization is a key event for blocking prodegenerative interactions (Figure 5G). Overall, our findings revealed that RBP4 blocks DR6 prodegenerative heterointeractions by forming an inactive 2:2 complex, necessitating the short IDR-dependency of DR6 dynamics.

### Increased Ca^2+^ directly exerts an agonistic propensity

Our identification of the antagonistic ligand RBP4 and its accompanying dynamic mechanism provides insights into the context and mechanism of DR6 signaling regulation. However, a fundamental question remained: What triggers DR6 signaling activation? This trigger could be inferred from the factors that interfere with DR6 homodimerization, which is primarily driven by IDR.

IDR-mediated protein interactions have emerged as a major factor in protein–protein interactions and biomolecule condensate formation.^60^ These phenomena are sensitive to nongenetic factors, such as pH, salt concentration, and temperature, enabling physicochemical cellular responses to the environmental factors.^61–63^ Notably, during axon pruning after TF deprivation, the initiation of the DR6-mediated catastrophic phase largely depends on the rupture of spheroid-like structures and the consequent release of Ca^2+^.^20^ Ca^2+^ serves as a universal second messenger crucial for neuronal synaptic activity and is tightly regulated in neurons. Extracellular Ca^2+^ dysregulation is causally implicated in many cell death and neurodegeneration processes and is a common feature of aging and neurodegenerative disorders.^64^ Notably, the mechanistic implication of extracellular Ca^2+^ has been mainly understood as a cellular “re-uptake” and activation of intracellular prodegenerative pathways, such as the calpain cascade and ER stress.^20^ Conversely, the direct contribution of extracellular Ca^2+^ to a cell death pathway remains poorly characterized.

Given the general properties of IDRs in salt sensitivity and the involvement of Ca^2+^ in DR6 signaling and neurodegeneration, we hypothesized that extracellular Ca^2+^ can directly trigger DR6 dissociation and signaling activation. To test this hypothesis, we determined the effect of Ca^2+^ on the dynamic oligomerization of DR6 ECD using size exclusion chromatography. On the elution profile resulting from the dynamic protein oligomerization, results revealed a direct interference with dynamic ECD oligomerization upon supplementation with 5 mM CaCl_2_, suggesting that DR6 ECD oligomerization directly responds to Ca^2+^ (Figure S6A and S6B). Next, we investigated the effect of increasing Ca^2+^ concentrations on cis interaction with APP and p75^NTR^. Aligning with the cognate link to DR6 dynamics revealed through RBP4, we found that Ca^2+^ directly promoted heterotypic interactions (Figure 5H). However, the effect of CaCl_2_ was independent of the asparagine/serine-rich region, suggesting that Ca^2+^ acts globally or at site independent of this IDR (Figure S6C and S6D). The direct effect of Ca^2+^ was evidenced by decreased interaction following treatment with membrane-impermeable Ca^2+^-specific chelator (2 mM BAPTA or 5 mM EGTA; Figure 5I). These findings suggest that Ca^2+^ acts as an agonistic stimulus, directly inhibiting DR6 ECD oligomerization and facilitating interactions with APP and p75^NTR^.

Seeking to prioritize the regulatory role of RBP4 and Ca^2+^, we measured the effect of sufficient RBP4 on increasing Ca^2+^ concentrations, given the constitutive RBP4 presence in peripheral organs. Interestingly, the presence of 0.5 mM RBP4 significantly attenuated the effect of Ca^2+^ (Figure 5J and 5K). Under intact physiological conditions, the extracellular Ca^2+^ concentration in the mouse brain is approximately 1.2 mM. In sensory neuron culture media, the concentration rapidly increases to approximately 2 mM after 24 h of NGF deprivation. Thus, RBP4 significantly, but incompletely, impeded the effect of prodegenerative concentration of extracellular calcium on DR6. Collectively, our findings reveal an agonistic action of Ca^2+^ on DR6 signaling, which is modulated by RBP4.

### RBP4 exerts a neuroprotective effect in DR6-dependent axon pruning and neuronal death models

We present the molecular landscape of DR6 signaling regulation in which RBP4 functions as a principal regulatory ligand that blocks the heterotypic prodegenerative interactions with APP and p75^NTR^. Next, we sought to expand the findings of the molecular assays and confirm that RBP4 serves as a functional ligand that antagonizes DR6 signaling and exerts a protective effect within the biological context. However, we encountered challenges in defining *in vivo* models owing to the direction of signaling antagonism and the possible contribution of altered retinol delivery in complex biological systems. Thus, we examined whether RBP4 exerts a protective effect in previously defined DR6-dependent axon pruning and neuronal death models (Figure 6A).

**Fig. 6.**
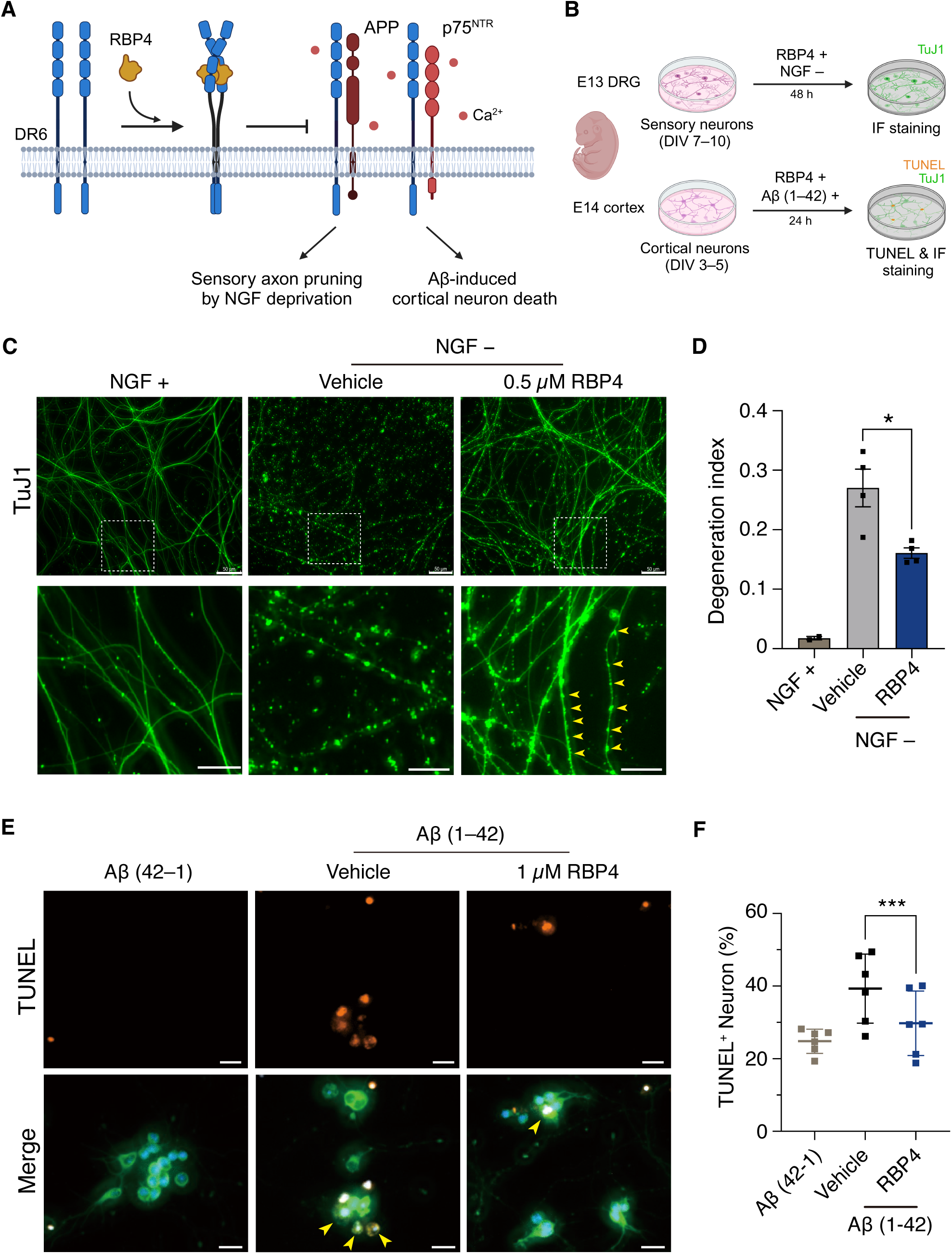
RBP4 prevents NGF-deprived sensory axon pruning and Aβ-induced cortical neuron death. (**A**) Scheme depicting the prodegenerative heterotypic interactions of DR6 and causally implicated biological processes. (**B**) Scheme depicting immunofluorescence staining (IF) of sensory axons cultured from embryonic dorsal root ganglion (DRG) after NGF deprivation and terminal deoxynucleotidyl transferase dUTP nick end labeling (TUNEL) staining of cortical neurons after treatment of oligomeric Aβ (1–42). (**C**) Representative image showing mouse sensory axon pruning after 48 hours of NGF deprivation and the protective effect of recombinant RBP4 (0.5 μM). The yellow arrow indicates the morphological feature of spheroid-like structure. Scale bars, 50 μm (upper) and 20 μm (lower). (**D**) Quantification of axonal degeneration by the ratio of fragmented axons. Data are presented as mean ± SD determined from four independent experiments. \**p*<0.05 by unpaired two-tailed t-test. (**E**) Representative image showing TuJ1-positive (green) mouse cortical neurons co-stained with TUNEL after 24 hours of treatment of oligomeric Aβ (1–42) or control reverse peptide (50 μg/ml). Neurons were TUNEL-stained and immunostained with TuJ1. The yellow arrow indicates the TUNEL^+^ neurons. Scale bars, 10 μm. (**F**) Quantification of TUNEL^+^ neurons. Data are presented as mean ± SD determined from six independent experiments counting at least 300 neuronal nuclei. \*\*\**p*<0.0005 by paired two-tailed t-test.

TF deprivation-induced axon pruning is one of the widely studied models crucially implicated in developmental sculpting, injury response, and neurodegenerative disorders. A recent study characterized the distinct phases of axon pruning in which the p75^NTR^-dependent latent phase drives the formation of a spheroid-like structure followed by a DR6-dependent catastrophic phase executing the caspase 6-dependent axon pruning process. During this process, DR6 signaling is critical for initiating the catastrophic phase, and signaling blockade produces an intact axonal morphology with a spheroid-like structure. Therefore, via qualitative and quantitative measurement of the axon pruning process, we can measure whether RBP4 blocks DR6 signaling and exerts a neuroprotective effect.

DR6 and RBP4 orthologs are only found in Vertebrata. Between humans and model organisms (*Mus musculus* and *Rattus norvegicus*), RBP4 and DR6 (excluding IDRs) exhibit high sequence homology (Figure S7A and S7B) and IDR properties (Figure S5D), supporting the conservation of the RBP4–DR6 axis. Thus, we tested the neuroprotective effect of murine RBP4 in the previously characterized mouse axon pruning and neuronal death models (Figure 6B).

Consistent with our previous findings, RBP4 exerted a prominent neuroprotective effect in NGF-deprived sensory axons with the formation of a spheroid-like structure (Figure 6C and 6D), indicating DR6 signaling blockade. Next, we tested whether RBP4 exerts a neuroprotective effect in oligomeric Aβ-treated cortical neurons in which the DR6–p75^NTR^ interaction is mechanistically implicated. Notably, neuronal death was suppressed by cotreatment with RBP4 and with oligomeric Aβ, suggesting a general neuroprotective effect of RBP4 (Figure 6E and 6F). Together with our finding on the protective effect in sodium arsenite-treated SH-SY5Y cells (Figure 2F), we found that RBP4 plays a major regulatory role in DR6-dependent cell death and neurodegenerative processes, highlighting the functional role of RBP4 as a DR6 antagonistic ligand.

## Discussion

The longstanding unsuccessful history of DR6 deorphanization studies, which mainly assumed the canonical death ligand-dependent model, has hindered a comprehensive understanding of the regulatory landscape of extrinsic regulation of cell death processes. In this study, through ePL and single-molecule analyses, we uncovered the distinct regulatory mechanism of DR6, as its antagonistic ligand RBP4 controls the autonomous prodegenerative hetero-interactions with APP and p75^NTR^ by forming an inactive 2:2 complex (Figure 7A).

**Fig. 7.**
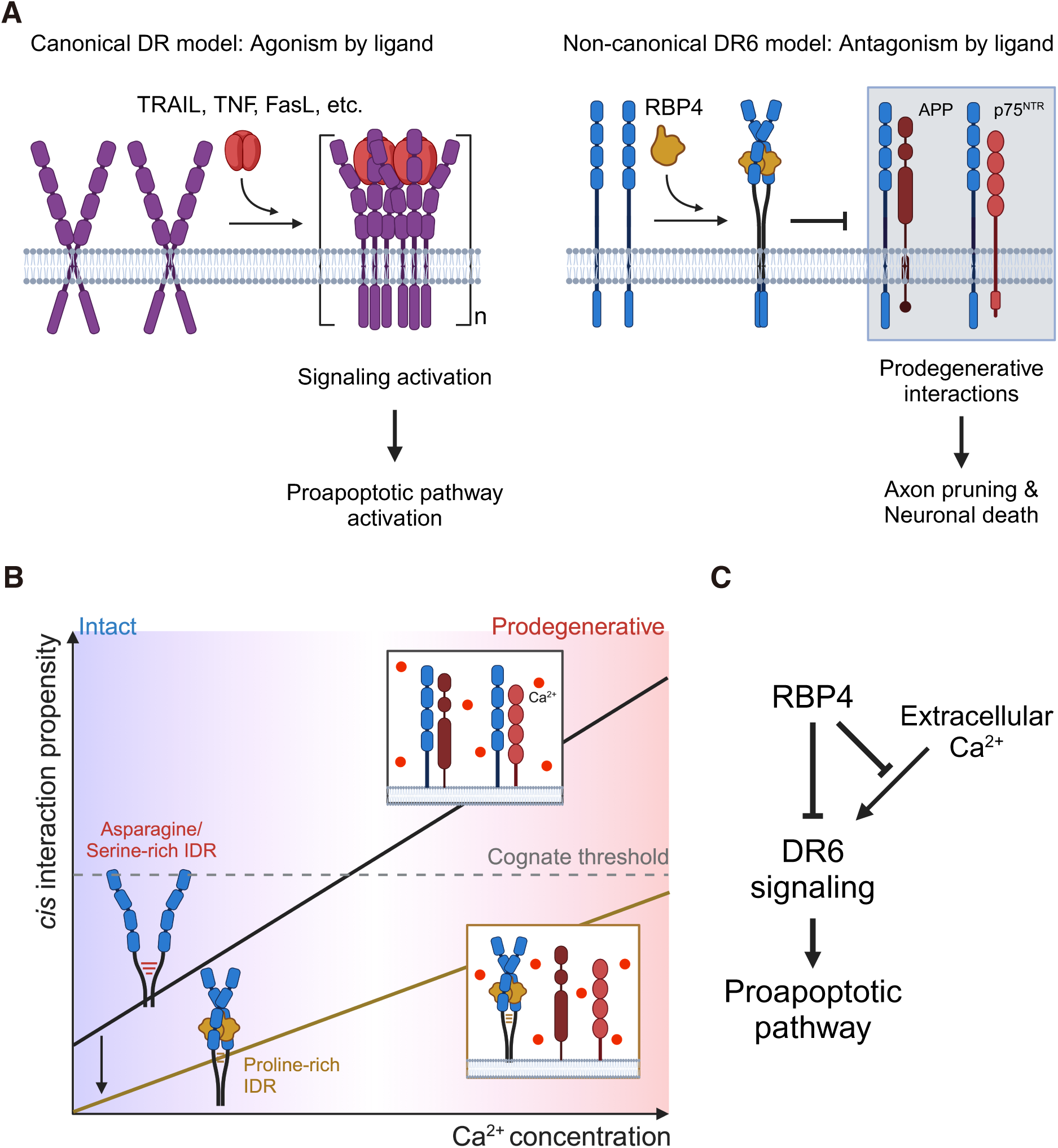
Scheme depicting the model of DR6 signaling regulation. (**A**) Scheme showing the comparison of ligand mechanisms between DRs and DR6. (**B**) Scheme depicting the model for DR6 signaling regulation by RBP4 and Ca^2+^. (**C**) Diagram showing DR6 regulation by RBP4.

The non-canonical death receptor mechanism could explain the distant sequence homology of DR6 from other DRs (Figure S8). Our unexpected findings on the direct regulatory role of RBP4 and concomitant molecular mechanism will deepen our understanding of the cellular program that potentially links vitamin A metabolism and cell death.

Understanding the ligand and molecular mechanism of DRs is of paramount importance to unwind the complex nature of biological processes and managing disease pathologies. Recently, therapeutic strategies that modulate DR signaling have been developed to block ligand interactions or receptor oligomerization or induce targeted receptor clustering in cancer cells. Among these strategies, several antibody drugs have been recently tested in clinical trials, highlighting the potency of DRs as druggable targets.^65–67^ DR6 is causally implicated in several pathogenic mechanisms, including autoimmune diseases, neuronal death in neurodegenerative disorders and tumor proliferation and metastasis. Our findings on the ligand and molecular mechanism reveal a novel avenue to modulate this signaling pathway and develop therapeutic options for various diseases.

RBP4 is expressed in several tissues and organs, such as adipose tissue, the brain, and the pancreas; however, liver-secreted holo-RBP4 is a major source of circulating RBP4 in normal physiology. The dissociation of RBP4 can lead to DR6 signaling activation. However, the dissociate-to-activate model of constitutively circulating retinol carriers alone does not fully provide temporal specificity to DR6 signaling activation. Instead, the temporal regulation of DR6 signaling can be achieved by modulating the calcium concentration, which dynamically ranges following the various prodegenerative biological contexts, including TF deprivation, cellular stress, and neuronal resting (Figure 7B). IDR-dependent DR6 dimerization sensitively and dynamically responds to Ca^2+^ (Figure S6A); however, these responses are almost linear, resulting in limited discrimination between physiological and degenerative concentrations (Figure 5H). Thus, a small protein that constitutively circulates and crosses the BBB could be used to prevent the uncontrolled Ca^2+^ response and modulate the threshold of the Ca^2+^ response along with additional spatial and biological contexts (Figure 5J). Because RBP4 constitutively circulates and Ca^2+^ serves as a global second messenger, RBP4-modulated Ca^2+^-responsive DR6 signaling could play a crucial role in fine-tuning the cellular stress responses to determine cell death (Figure 7C).

Vitamin A deficiency (VAD) remains a global health issue causing immune impairment, blindness, neurodegeneration, and consequent death.^68^ It limits the secretion of holo-RBP4 and decreases circulating RBP4 levels.^44^ Currently, the mechanistic bases of the pathological consequences of VAD have mainly been identified as impaired retinol delivery and cellular retinol responses. However, the detailed mechanism leading to ocular phenotypes, immune impairment, and neurodegeneration remains incompletely understood. As DR6 signaling regulates T and B cell function and the pruning of retinal ganglion cell axons, VAD-induced RBP4 decline might contribute to DR6 dysregulation and the pathogenic mechanism of VAD. Moreover, RBP4 decline and retinal degeneration are the universal features of DR6-associated neurodegenerative disorders, including AD, ALS, and MS.^69–72^ As the blockade of DR6 signaling in rodent models of ALS and MS exerted a prominent protective effect, RBP4 decline might also contribute to DR6 dysregulation, representing an initiative process of sporadic neurodegenerative consequences accompanying retinal degeneration. Thus, further *in vivo* studies are required to evaluate the pathogenic contribution of the RBP4–DR6 axis in VAD and sporadic neurodegenerative disorders.

In this study, we explicitly investigated the regulatory role of RBP4, which was identified from the secretome profiled by TurboID-DR6 in SH-SY5Y cells. The results of secretome profiling suggest additional aspects of the DR6 signaling regulatory molecule network. For instance, PRR4 and LACRT, eye-specific proteins with cognate protective roles^73,74^, were highly and predominantly enriched by TurboID-DR6. They could be the functional ligands of DR6 that specifically regulate eye-specific DR6 signaling regulation and cellular function.

Additionally, we observed that the ePL pattern was cell type-specific. As DR6 signaling is also implicated in endothelial cell necroptosis and immune responses, the cell type-specific ePL pattern might inform other possible regulatory landscapes of DR6. Thus, the application of ePL could help expand our understanding of complex trans-signaling regulatory mechanisms and cis-responsive processing of transmembrane receptors.

In conclusion, we characterized the regulatory mechanism of DR6, the last orphan member of the DR family. Unexpectedly revealing dynamic DR6 regulation by RBP4 and Ca^2+^ has advanced our comprehension of extrinsic regulation of cell death processes with implications for developing therapeutic strategies for many disease pathologies.

### Limitations of the study

In this study, we mainly focused on establishing the molecular mechanism of orphan DR6 regulation through RBP4 using molecular and biophysical tools and information. Our findings offer insights into the regulatory mechanism of DR6. Although we studied the protective effect of RBP4 in DR6-dependent models, further *in vivo* studies are required to evaluate the role of RBP4 in DR6 signaling regulation and associated phenotypes.

Additionally, although ePL offers other possible ligand candidates, such as PRR4 or LACRT, we predominantly focused on RBP4 as a DR6 ligand. Owing to the proximal limitation of N-terminal TurboID, the ligand candidates interacting with other binding sites or IDRs can be discovered through other methods, necessitating additional ligand search studies.

## Acknowledgments

We thank J.K, J.I, and the members of WARC, especially B.C, H.C, and J.J for discussions; H.R, G.Y, and D.K for the gift of Sodium bicarbonate, TurboID plasmid, and Streptavidin-HRP; G.J and C.L for the use of Alphafold; Y.M for the preparation of mouse sensory neuron culture. This work was supported by the National Research Foundation of Korea (NRF) grant funded by the Korea government (MSIT). (2021R1A2C1009107)

## Author contributions

T.K designed, performed, and interpreted most of the experiments. T.N.C.N and M.-S.K performed and interpreted LC-MS^2^ analysis. J.J performed and interpreted the single-molecule photobleaching assays and K.K performed and interpreted the single-molecule DR6 tracking experiments, both under the guidance of W.J.L and D.S. T.K wrote the manuscript with input from coauthors. Y.-S.L provided critical advice. Y.-I.L supervised the project. All the authors critically reviewed the manuscript.

## Supplementary information

Figures S1–S8

Table S1. Excel file containing the complete list of biotinylated protein, related to Figure 1

Table S2. Excel file containing the list of crosslinked residues between RBP4 and DR6, related to Figure 2

Video S1–S3. Tracking of QD-labeled DR6 and diffusive state analysis, related to Figure 4.

