## Supplementary information for "RBP4 functions as an antagonistic ligand of orphan DR6"

### Materials and Methods

#### Mice

Pregnant (E13–14) ICR mice were purchased from Daehan Biolink Co., Ltd. (Umsung, Chungbuk, South Korea). Animals were sacrificed by the physical method of cervical dislocation, and the embryos were collected in DPBS, and cortices or dorsal root ganglion (DRG) were dissected for primary culture. All mice were handled following the animal care standards outlined in the Korean National Institute of Health Guide for the Care and Use of Experimental Animals and the experiments were approved by the Daegu Gyeongbuk Institute of Science and Technology (DGIST) Administrative Panel on Laboratory Animal Care. (DGIST-IACUC-22072708-0000)

#### Cell lines

SH-SY5Y and HEK-293T cell lines were gifted from Cheil. Moon (DGIST). Hs683 cell line was purchased from Korean Cell Line Bank (South Korea). Cells were maintained in cell culture media (DMEM (Welgene) supplemented with 10% fetal bovine serum (Welgene) and 1% penicillin-streptomycin (Welgene)) in a humidified atmosphere of 5% $CO_2$ at 37 °C.

#### Plasmid construction

Plasmids were selected for the purpose of each experiment. For protein expression and purification in E.coli system, pET21a (mouse and human RBP4; NdeI–XhoI) and pET28a (mouse RBP4; NcoI–XhoI) were used. For mammalian expression and purification, genes of interest were inserted into the pLexm plasmid between EcoRI–XhoI sites. For mammalian expression under the control of CMV promoter, genes of interest were subcloned into pLenti-C-Myc-DDK-P2A-Puro cloned with human Tnfrsf21 (DR6) plasmid purchased from Origene (#RC205864L3) between AsiSI–MluI restriction sites. For mammalian expression under the control of HSV TK promoter, genes of interest were subcloned into pRL-TK vector purchased from Promega between NheI–NotI restriction sites.

#### Recombinant proteins

Recombinant mouse or human RBP4 was purchased from R&D systems or purified by oxidative refolding from E.coli inclusion body (IB). Purchased recombinant RBP4 was incubated with 10 times molar mass of All-trans retinoic acid (ATRA; Sigma-Aldrich) in PBS for 4 h at room temperature (RT) and ultrafiltrated by Amicon Ultra 10K 0.5 ml (Millipore) to remove unbound ATRA. Recombinant human DR6 ECD was purchased from Sino Biological. Recombinant RBP4-SNAP and DR6-Halo were expressed in SH-SY5Y transiently transfected by lipofectamine 2000 (Thermo Fisher scientific) and purified by Ni-INDIGO agarose beads (Cube Biotech) in cultured media.

**Fluorescence microscopy**

Fluorescence microscopy of mouse sensory and cortical neurons was performed by Axio Observer Z1 (Carl Zeiss) with 4× or 10× magnification.

For single molecular imaging, Total Internal Reflection Fluorescence (TIRF) microscopy system including an inverted microscope (Nikon, ECLPSE Ti2-E), a motorized stage (TI2-S-SE-E), an electron multiplying charge-coupled device (EM CCD, Andor, iXorn Ultra 897), a spectroscopic detector (Andor, Newton DU-971), a perfect focus system (PFS, TI2-N-ND-P), and laser light sources (Nikon, LU-N4 Laser Unit, 488 nm, 640 nm wavelength) was used. Images were captured with a 100× oil-immersion objective lens (Nikon, 1.49 NA, CFI SR HP Apochromat TIRF).

**DR6 ligand candidate enrichment using proximity labeling**

DNA sequence encoding TurboID-DR6 or control was inserted into pLenti-C-Myc-DDK-P2A-Puro. For a small-scale enrichment for WB, SH-SY5Y cells were seeded in a 12-well plate at $2\times10^5$ density. The next day, 500 ng of plasmid encoding TurboID-DR6 or negative control was transfected using lipofectamine 2000. After 24–48 hours incubation, the media was removed and changed to biotinylation media (cell culture media with 250 μM biotin (Sigma-Aldrich) and 1 mM ATP (Sigma-Aldrich)). Cells were incubated with biotinylation media for 24 hours, and the media was collected, filtrated by 0.45 μm syringe filter, and concentrated by Amicon Ultra 10K 0.5 ml at 4 °C. Concentrated media was sequentially diluted and concentrated with pre-chilled PBS to reach 5,000–10,000 times dilution. Diluted media was then incubated overnight with Pierce Streptavidin Magnetic Beads (Thermo Fisher Scientific), 1× Halt protease inhibitor cocktail (Thermo Fisher Scientific), and 5 mM EDTA (Thermo Fisher Scientific) at 4 °C. Beads were collected by magnetic rack and washed twice by radioimmunoprecipitation assay buffer (RIPA; Biosesang), once by 1 M KCl (Sigma-Aldrich), briefly once by 0.1 M $Na_2CO_3$ (Sigma-Aldrich), briefly once by 2M urea (Sigma-Aldrich) in 10 mM Tris-HCl (pH 8.0; Biosesang), and twice by RIPA.[75] After washing, proteins bound to beads were eluted by directly boiling the beads in 6× sample buffer (0.33 M Tris-HCl (pH 6.8; Biosesang), 34% (v/v) glycerol (Sigma-Aldrich), 10% (w/v) sodium dodecyl sulfate (SDS; Sigma-Aldrich),

0.09% (w/v) DL-Dithiothreitol (DTT; Goldbio), 0.12% (w/v) bromophenol blue (Sigma-Aldrich) with 100 mM DTT and 2 mM biotin at 95 °C for 10 min. The eluted proteins in the sample buffer were analyzed by WB.

For large-scale enrichment for LC-$MS^2$, SH-SY5Y cells were transiently transfected TurboID-DR6 or TurboID-EGFP/DR6 by lipofectamine 2000. After 48 hours, cells were treated with 2 μg/ml puromycin (Gibco) for 2 weeks to generate a stable cell line. Stable cells were seeded to a 175T flask. The next day, cells were incubated with 30 ml biotinylation media. After 24 hours incubation, media was collected, filtrated by Steriflip 50 ml vacuum filter (Sigma-Aldrich), and concentrated by Amicon Ultra 10K 15 ml at 4 °C. Concentrated media was sequentially diluted and concentrated with pre-chilled PBS to reach 5,000-10,000 times dilution. Diluted media was then incubated overnight with Pierce High Capacity Streptavidin Agarose Beads (Thermo Fischer Scientific), 1× Halt protease inhibitor cocktail, and 5 mM EDTA at 4 °C. The beads were washed thrice by RIPA, briefly thrice by Tris-buffered saline (TBS; Biosesang), and resuspended in 1 ml of 0.1 M $NaHCO_3$ (Sigma-Aldrich) and 2 M urea (pH 7.76) and moved to a microcentrifuge tube. The beads were collected and washed twice by RIPA. The proteins were then eluted by directly boiling the beads in 6× sample buffer with 100 mM DTT and 2 mM biotin at 95 °C for 10 min. The eluted proteins in the sample buffer were separated by SDS-PAGE and stained by Coomassie blue. The visible protein bands from the TurboID-DR6 sample and size-matched bands from the TurboID-EGFP/DR6 sample were cut with a clean knife, and gel pieces were stored in distilled-deionized water (DIW) at 4 °C for in-gel digestion processing.

**Identification of biotinylated proteins using LC-$MS^2$**

The gel cuts were destained in 40% acetonitrile (J.T Bakers) with 25 mM ammonium bicarbonate (ABC; Sigma-Aldrich) and 10 mM DTT prepared in DIW for 3 cycles of 10 min. Proteins were then alkylated by 10 mM iodoacetamide (Sigma-Aldrich) for 30 min protected from light. After washing with 40% acetonitrile with 25 mM ABC, the gel cuts were dehydrated by acetonitrile. Proteins were in-gel digested by trypsin in 50 mM ABC overnight and peptides were extracted from the gel by 40% acetonitrile with 0.1% formic acid, followed by desalting using a C18 disk (Empore™; 3M).

Biotinylated peptides were analyzed on a Q-Exactive Orbitrap mass spectrometer (Thermo Scientific) coupled with a liquid chromatography system (EASY-nLC; Thermo Scientific). Peptides were loaded on a trap column (75 μm × 2 cm; C18; Acclaim PepMap100; Thermo Fisher Scientific) and subsequently separated on an analytical column (75 μm × 50 cm; C18;

PepMap RSLC; Thermo Fisher Scientific) at a flow rate of 250 nl/min. A 2-hour gradient started with 5% solution B (98% acetonitrile, 2% water, and 0.1% formic acid) and stabilized for 3 min followed by a linear increase to 27% B for 97 min, then ramped to 90% B for 3 min and maintained at 90% for 3 min before dropping to 5% B for 5 min and equilibrated at 5% B for 5 min. Data were acquired on the Orbitrap with the following parameters: MS1 scan range was from 300 to 2000 m/z, resolution of 70,000, maximum ion injection time of 20 ms, automatic gain control (AGC) target of $10^6$. $MS^2$ spectra were produced by high energy collision dissociation with 27% normalized energy and scanned at a resolution of 35,000 and a maximum ion injection time of 50 ms.

Raw tandem mass spectra of biotinylated peptides were processed by MaxQuant (v1.6.17.0; PMID: 27809316) and searched against a human proteome database obtained from Uniprot. Carbamidomethylation on cysteine and biotinylation on lysine (+226.078 Da) were set for variable modifications while oxidation on methionine and acetylation on protein N-termini were for fixed modifications. False discovery rates (FDRs) were 1% for both peptides and proteins. Mass tolerance of the first search and main search was 20 ppm and 4.5 ppm, respectively. The minimum peptide length was 7 with a maximum of 2 mis-cleavages by trypsin.

**Affinity purification of antibodies in media**

To induce the biotinylation of control mouse IgG or DR6 antibody (Santa Cruz Biotechnology), $2\times10^5$ SH-SY5Y cells were seeded in a 12-well plate and transfected with 500 ng of TurboID-DR6. The next day, the media was removed, and cells were incubated with 10 μg/ml antibodies in biotinylation media for 24 hours. After incubation, media was collected, filtrated, and incubated with Protein A/G Plus-Agarose (Santa Cruz Biotechnology) overnight. The beads were washed thrice with Pierce IP lysis buffer (Thermo Fischer Scientific) and eluted by directly boiling them in the 6× sample buffer with 10 mM DTT. For collecting the antibodies by streptavidin beads, sample preparation was performed by following the same protocol with biotinylated protein enrichment in the media. The eluted antibodies were then analyzed by WB.

**Cell death assay**

2 M sodium (meta)arsenite (Sigma-Aldrich) solution prepared in DIW was filtrated and stored at −20 °C. $2\times10^5$ SH-SY5Y cells were seeded in 12-well plate and transiently transfected with 500 ng plasmid (GFP, DR6, or TurboID-DR6) by lipofectamine 2000. The next day, the media was changed to cell culture media containing 300 or 350 μM sodium arsenite. After 24 hours, cells were collected by centrifugation and resuspended in DPBS (Welgene). Cells were stained

with trypan blue, and permeabilized cells were automatedly counted by Countess II FL (Thermo Scientific). The number of permeabilized cells per total cells was used to calculate the cell death ratio.

**Disease association and gene ontology analysis**

The Database for Annotation, Visualization, and Integrated Discovery (DAVID; v2023q4; https://davidbioinformatics.nih.gov) was used for gene ontology analysis for secretory proteins identified by LC-$MS^2$ from TurboID-DR6 and TurboID-EGFP/DR6 cultured media.[76,77] The complete protein list was uploaded under the heading of UP_KW_CELLULAR_COMPONENT. The proteins annotated to be 'secreted' were then uploaded to DAVID under the heading of GAD_DISEASE and GOTERM_BP_DIRECT. To check the biased protein enrichment from neuroblastoma cell line, the list of secretory proteins enriched in TurboID-EGFP/DR6 and TurboID-DR6 was uploaded as background.

**Network analysis**

The list of secretory proteins enriched by TurboID-DR6 was uploaded to NetworkAnalyst (https://www.networkanalyst.ca).[78] Zero-order Protein-Protein Interaction network was generated using IMEx interactome database.

**Western blot**

The proteins were separated by 4–20% Novex Tris-Glycine Mini Protein Gels (Thermo Fisher Scientific) and transferred to the nitrocellulose membrane (Amersham Pharmacia Biotech). Membranes were blocked by 5% skimmed milk (KisanBio) in TBST (TBS with 0.1% (v/v) Tween-20 (Amresco)) with rocking at RT for 1 hour. The membrane was incubated with primary antibody or streptavidin-HRP (1:2000, Cell Signaling) in 3% BSA (Lee Biosolutions) in TBST for 1 hour with rocking at RT. For protein detection using streptavidin-HRP, the membranes were washed 5–6 times by TBST for 5 min with orbital shaking at RT. For protein detection by antibodies, the membranes were washed 3 times by TBST for 5 min with orbital shaking at RT. Membranes were then incubated with HRP-conjugated anti-mouse, rabbit, or goat antibodies (1:5000) in 5% skimmed milk in TBST for 1 hour with rocking at RT and washed thrice by TBST for 5 min with orbital shaking at RT. The HRP signals were visualized by chemiluminescence (Immobilon Western HRP substrate; Sigma-Aldrich) with ImageQuant LAS-4000 (GE healthcare) or Amersham Imager 680 (GE healthcare).

The primary antibodies used were as follows: anti-beta-amyloid (1:2000, Proteintech), anti-RBP4 (1:2000, Proteintech), anti-IGFBP2 (1:2000, Proteintech), anti-clusterin (1:2000, Proteintech), anti-FSTL1 (1:2000, Proteintech), anti-TGFB1 LAP (2 µg/ml, R&D systems), anti-GAPDH (1:2000, Santa Cruz Biotechnology), anti-DR6 (1:2000, Santa Cruz Biotechnology), and anti-HIS tag-HRP (1:2000, Santa Cruz Biotechnology).

**Oxidative refolding and Purification of recombinant RBP4**

Recombinant RBP4 was expressed and purified as previously described with modifications.[79] Briefly, cDNA encoding mouse and Human RBP4 without signal peptide was subcloned into pET21a vector, and plasmids were transformed into Rosetta plysS competent cells. For expression, E.coli was first pre-cultured in 3 ml LB Broth (KisanBio) with 100 µg/ml ampicillin overnight. The next day, 2 ml pre-cultured cells were inoculated into 1 L LB Broth and incubated at 37 °C with rotation. When OD600 reached 0.6–0.8, 1 M Isopropyl-ß-D-thiogalactopyranoside (IPTG) was added to make the final 1 mM concentration, and cells were incubated overnight. The next day, cells were collected by centrifuge at 3,000g for 15 min and washed with 0.9% NaCl (Sigma-Aldrich). Cell pellets were resuspended in lysis buffer (50 mM Tris-HCl (pH 7.5), 2 mM ethylene-diamine-tetraacetic acid (EDTA), 2 mM Phenylmethylsulfonyl fluoride (PMSF), and 0.1% triton X-100 (Sigma-Aldrich)) and sonicated. Cell lysates were then centrifuged at 12,000g for 15 min, and the pelleted IB was aliquoted, stored at −20 °C, and used for RBP4 oxidative refolding.

RBP4 in the IB was solubilized in a solubilization buffer composed of 2 mM Tris-HCl (pH 9.0; Biosesang), 5 M guanidium chloride (Tokyo Chemical Industry), and 10 mM DTT with overnight vigorous stirring. (25 ml solubilization buffer for 250 ml cultured E.coli) The next day, the translucent IB solution was centrifuged at 12,000g for 15 min, and the supernatant was collected and pre-chilled on ice. For 25 ml IB solution, 100 ml refolding buffer (25 mM Tris-HCl (pH 9.0), 0.3 mM Cysteine (Sigma-Aldrich), 3 mM Cystine (Sigma-Aldrich), 1 mM EDTA (Biosesang)) was prepared by degassing through helium sparging for 20 min on ice. Before mixing with solubilized IB, 3.2 ml of 15 mM ATRA prepared in ethanol was added to the refolding buffer with vigorous stirring. The solubilized IB was then rapidly mixed with refolding buffer and vigorously stirred for 5 hours at 4 °C protected from light. After refolding, the solution was centrifuged at 12,000g for 15 min at 4 °C. Supernatant was collected and, if necessary, flew through an empty Pierce Centrifuge Column (Thermo Fischer Scientific) by gravity flow to remove the precipitates. Refolded proteins were then extensively dialyzed against a buffer composed of 20 mM Tris-HCl (pH 9.0) and 150 mM NaCl using a 15–30 ml

Slide-A-Lyzer dialysis cassette (Thermo Fischer Scientific) protected from light until reaching 10,000-100,000 times dilution.

The dialyzed solution was incubated with pre-washed Ni-INDIGO agarose resin without imidazole (Sigma-Aldrich) supplement with overnight agitation at 4 °C. Beads were collected in Pierce Centrifuge Column by gravity flow, washed with 30 ml of 10 mM imidazole in dialysate, and eluted in 5 ml of 500 mM imidazole in dialysate. Protein eluent was briefly concentrated to reach 1 ml by Amicon Ultra 10K 0.5 ml with 10,000g centrifuge and loaded onto a Superdex 75 10/300 GL (Cytiva) gel filtration column filled with 20 mM Tri-HCl (pH 9.0), 150 mM NaCl, and 1 mM Tris(2-carboxyethyl)phosphine hydrochloride (TCEP; Sigma-Aldrich). UV absorption at 330 nm was monitored for Holo-RBP4. The collected Holo-RBP4 was concentrated, and the buffer was exchanged with assay buffer (HBSS (w.o. $CaCl_2$ and $MgCl_2$; Welgene) with 20 mM HEPES (Sigma-Aldrich; pH 7.4)). The protein concentration was determined by Pierce BCA Protein Assay Kit (Thermo Fischer Scientific), and 200–250 μM recombinant RBP4 and flow through (vehicle control) were snap-frozen by liquid nitrogen and stored at −80 °C.

**Surface plasmon resonance (SPR) analysis**

SPR was performed by SR7500DC (Reichert). Recombinant RBP4 was immobilized in a Polyethylene glycol (PEG)-coated sensor chip using the amine coupling following the manufacturer's instruction. The device was filled with running buffer (1 mg/ml PEG-8000 (Sigma-Aldrich) in PBS). The kinetic assay was performed by an increasing concentration of recombinant hDR6 ECD bound on the two flow cells (RBP4-bound active and unbound reference cell). The signal from the active cell subtracting the signal from the reference cell was measured.

**Identification of DR6-RBP4 crosslinked sites by LC-MS$^2$ (XL-MS)**

Recombinant hDR6 ECD and RBP4 prepared in assay buffer were mixed at 10 μM concentration and incubated for 3 hours at 4 °C. Then, an equal volume of 1 mM BS$^3$ (Thermo Fischer Scientific) freshly prepared in the same buffer was mixed to make a final concentration of 0.5 mM, and the mixture was incubated for 2 hours at 4 °C. The crosslinking reaction was quenched by adding 50 mM Tris-HCl (pH 7.5). The crosslinked mixture was then deglycosylated by Protein Deglycosylation Mix II (New England Biolabs) following the manufacturer's instructions. The buffer was briefly changed to assay buffer and mixed with WB sample buffer. The proteins were separated by SDS-PAGE using NuPAGE 3–8% Tris-Acetate gel (Thermo

Fischer Scientific), and the gel was stained by Coomassie blue. The crosslinked protein complex was identified by comparing it with WB data run on the same gel. The gel pieces were cut by a clean knife and stored in DIW for in-gel digestion processing.

Crosslinked proteins were also processed by the in-gel digestion method described above. Crosslinked peptides were analyzed on a Q-Exactive Orbitrap mass spectrometer (Thermo Scientific) coupled with Vanquish Neo (Thermo Scientific). Peptides were eluted with a flow rate of 300 nl/min along the trap column and analytical columns as mentioned above. The gradient started with 5% solution B (0.1% formic acid in acetonitrile) and 95% solution A (0.1% formic acid in HPLC water) for 2 min, followed by 30% B for 100 min and inclined to 90% B for 10 min prior to 5% B for 5 min and equilibrated at 5% B for 3 min. Peptides were ionized and analyzed on the Q Exactive Orbitrap (Thermo Scientific). MS1 scans were acquired with the following parameters: scan range from 350 to 1650 m/z, resolution of 35 000, maximum IT of 20 ms, and ACG target of $10^6$. Data were acquired under DDA mode and ions with charge state from 2 to 7 and >8 were selected for fragmentation at 27% collision energy and resolution of 17 500, maximum IT of 50 ms, and AGC target of $10^5$.

Raw tandem mass spectra of the peptides were processed by MaxQuant (v2.3.1.0). The crosslinked peptides were searched against the sequences and reversed sequences (as decoys) using Xisearch (v1.7.6.7). The following parameters were applied for the search: MS1 accuracy = 6 ppm; MS2 accuracy = 20 ppm; enzyme = trypsin (with full tryptic specificity) allowing up to two missed cleavages; crosslinker = $BS^3$ with an assumed reaction specificity for lysine, serine, threonine, tyrosine and protein N termini; fixed modifications = carbamidomethylation on cysteine; variable modifications = oxidation on methionine, hydrolyzed/aminolyzed $BS^3$ from reaction with ammonia or water on a free crosslinker end. The identified candidates were filtered to 1% FDR on link level using XiFDR (v2.1.5.5).

**Protein docking and model selection**

Protein docking simulation was performed through the Cluspro 2.0 web server, which uses PIPER algorithm to find the paired interactions and generates the docking models ranked by the interaction energy, stability, and the size of clustered models (https://cluspro.bu.edu/).[80] The crystal structure of DR6 CRD (PDB #3U3P) and Holo-RBP4 (PDB #5NU7) deposited in PDB was uploaded to Cluspro 2.0. The structure of DR6 CRD was defined as a receptor, and RBP4 was defined as a ligand. Crosslinked sites identified by XL-MS analysis were used as a restraint. If one residue was crosslinked with multiple residues, the residue with the highest score calculated by XiFDR was selected. DR6-RBP4 residue pairs were selected as follows;

I51(K46)-S64, K159-K168, T175-T146, and T132-E19. Given the length of $BS^3$ and lysine residues and the distance from unstructured K46 to structured I51 of DR6, the minimum ('dmin') length of restraints was set to 6, and the maximum ('dmax') length was set to 30, 40, 14, and 14, respectively. The restraint information was written in JSON file and uploaded to Cluspro 2.0.

**Structural prediction through Alphafold**

The structure of mouse and human RBP4 and CRD of human DRs were predicted by Alphafold installed and run on a Linux machine.[81] (https://github.com/google-deepmind/alphafold)
The prediction of a 2:2 binding structure using Alphafold3 was performed by Alphafold3 web server (https://golgi.sandbox.google.com).[82] Protein sequences encoding DR6 ECD (42-349) and RBP4 were uploaded.

**Bioluminescence Resonance Energy Transfer (NanoBRET) assay**

For 5-TMR labeling of RBP4, 500 μg purified recombinant human RBP4 was concentrated and buffer was exchanged to 50 mM borate buffer (pH 8.8; Thermo Scientific) by Amicon Ultra 10K 0.5 ml. 1 mg of 5-Carboxy-tetramethylrhodamine, *N*-succinimidyl ester powder (5-TMR, SE; GLPBIO) was directly resolved in 2.5 mg/ml RBP4 in 50 mM borate buffer. The labeling reaction was proceeded by incubating at RT for 1 hour. After labeling, the reaction was quenched by 50 mM Tris-HCl (pH 8.5) supplement. Unreacted dyes were removed and the buffer was changed to assay buffer using 0.5 ml Amicon ultra 10K until the dilution reached 10,000 times. 5-TMR-RBP4 was then stored at 4 °C protected from light, and used for Nano-BRET assay.
HEK-293T cells were seeded in 96-well white plates at $2\times10^4$ density. The next day, cells were transfected with 50 ng plasmid encoding Nanoluciferase-DR6 WT or mutants. After 48 hours, cells were incubated with serially diluted 5-TMR-RBP4 in assay buffer and 200× Vivazine luciferase substrate (Promega). After 60 min incubation at 37 °C incubator, dual color luminescence was detected by Infinite M200 pro (Tecan). The BRET ratio was calculated by subtracting GREEN1 from BLUE1 channel intensity. The average of the BRET ratio from 4 wells at each 5-TMR-RBP4 concentration was compared to the untreated control.

**Luciferase complementation assay (NanoBit)**

Owing to the high endocytotic activity of DR6 and its binding partners, APP[83] and $p75^{NTR}$, [84] the effect of extracellular RBP4 was detectable only in specific experimental conditions. $2\times10^4$

HEK-293T cells were seeded in a 96-well white plate. After 16–24 hours from seeding, plasmids encoding DR6 (1–370) linked to SmBit through GGGGS linker and APP751 (1–703) or $p75^{NTR}$ (1–272) linked to LgBit through GGGGS linker under HSV TK promoter (pRL-TK) were transfected using lipofectamine 2000 at 50 ng each. After 18–22 hours post-transfection, media was removed, and cells were incubated with 50 μl assay buffer with 500x Vivazine luciferase substrate. After 1 hour incubation at 37 °C incubator, the luminescence signal was measured until stabilized (generally $4^{th}$ detection) and used as a basal signal. 50 μl of the same buffer containing 1 μM RBP4 or vehicle control was added to each well and incubated for 30 min at a 37 °C incubator. The stabilized luminescence signal was divided by the basal signal from each well. The relative signals from the basal level at each condition were obtained by the average of signals from 4 replicates, and the ratio to the signal from the vehicle control was calculated.

The effect of $Ca^{2+}$ on NanoBit assay requires substantial gene expression in the cells. $2\times10^4$ HEK-293T cells were seeded in a 96-well white plate. After 16–24 hours from seeding, plasmids encoding DR6 (1–414) linked to LgBit and APP751 N740A (endocytosis inhibited[85]) or $p75^{NTR}$ (1–343) linked to SmBit under CMV promoter (pLenti-C-Myc-DDK-P2A-Puro) were transfected using lipofectamine 2000 at 10 ng each. After 18–22 hours post-transfection, media was removed, and cells were incubated with 50 μl assay buffer with 500x Vivazine luciferase substrate. After 1 hour incubation at 37 °C incubator, the luminescence signal was measured until stabilized and used as a basal signal. 50 μl of 2× of the given concentration of $CaCl_2$ in the same buffer with Vivazine was then added to each well and incubated for 30 min at a 37 °C incubator. The stabilized luminescence signal was divided by the basal signal from each well. The relative signal from the basal level at each condition was obtained by the average of signals from 4 replicates, and the ratio to the signal from the control was calculated.

For measuring the effect of $Ca^{2+}$-specific chelators, BAPTA and EGTA, HEK-293T transiently transfected with the plasmids encoding DR6 (1–414) linked to LgBit and APP N740A or $p75^{NTR}$ (1–343) linked to SmBit under CMV promoter (pLenti-C-Myc-DDK-P2A-Puro) were incubated with 50 μl assay buffer with 2 mM $CaCl_2$ and 500x Vivazine luciferase substrate. After 1 hour incubation at 37 °C incubator, the luminescence signal was measured until stabilized and used as a basal signal. 50 μl of 2× $Ca^{2+}$ chelator (10 mM EGTA or 4 mM BAPTA) in the same buffer was then added to each well and incubated for 10 and 30 min at a 37 °C incubator. The stabilized luminescence signal was divided by the basal signal from each well. The relative signal from the basal level at each condition was obtained by the average of signals from 4 replicates, and the ratio to the signal from the control was calculated.

**NGF deprivation in mouse sensory neuron culture**

DRG from 12–15 E13 ICR embryos were dissected and collected in DPBS. DRG were then dissociated in 0.25% trypsin in DPBS by adding 2.5% trypsin solution (Hyclone) for 15 min at 37 °C water bath. Cells were then briefly triturated by mild pipetting. The digestion was stopped by adding an equal volume of cell culture media. Dissociated cells were collected by centrifugation at 440g, resuspended in sensory neuron culture media (Neurobasal media (Thermo Fischer Scientific) with 50 ng/ml NGF (Alomone Labs), 2% B-27 (Thermo Fischer Scientific), 1× Glutamax (Thermo Fischer Scientific), and 1% penicillin-streptomycin), and passed through a 40 μm cell strainer. Viable cells were automatedly counted by Countess II FL (Thermo Scientific) with trypan blue staining.

For coating cell culture slides, Flux 2-well cell culture slides (SPL Life Science) were incubated with 50 μg/ml Poly-D-lysine (PDL; Sigma-Aldrich) in DPBS overnight at 37 °C incubator and washed thrice with DBPS. The slides were then incubated with 2 μg/ml laminin (Sigma-Aldrich) in DPBS for 4 hours at 37 °C and washed thrice with DPBS before cell seeding.

Cells were seeded at $1.5\times10^5$ density in 500 μl sensory neuron culture media and incubated at 37 °C incubator. After 24 hours, 500 μl of 2× neuron purification media was added to the cells to make a 1× final concentration (sensory neuron culture media with 0.5 μM cytarabine (Sigma-Aldrich), 20 μM floxuridine (Abcam), 20 μM uridine (Sigma-Aldrich)). After 72 hours, 660 μl of media was replaced with fresh neuron culture media. After an additional 48 hours, 500 μl of media was replaced with fresh neuron culture media. The purified neurons were then used for NGF deprivation.

When DIV reached 8-10, neurons were washed twice with sensory neuron culture media without NGF. Neurons were then incubated with 1 ml NGF deprivation media (sensory neuron culture media without NGF and with 1 μg/ml anti-NGF antibody (Alomone Labs)) for 48 hours. Recombinant RBP4 or vehicle control was treated in NGF deprivation media.

Mouse sensory neurons were briefly washed with PBS and fixed by 4% paraformaldehyde (PFA; Biosesang) in PBS for 10-15 min at RT. Cells were washed thrice with PBS and blocked by blocking solution (10% donkey serum (v/v), 0.3% triton X-100 (v/v), and 0.1% BSA (w/v) in PBS) for 1 hour at RT. Cells were then incubated with anti-TuJ1 antibody (1:300; Abcam) in 1% donkey serum, 0.3% triton X-100, and 0.1% BSA in PBS overnight at 4 °C. Cells were washed thrice with PBS and incubated with 1% donkey serum and 0.1% BSA in PBS for 30 min at RT. Cells were then incubated with Alexa Fluor 488-conjugated anti-mouse antibody (1:300; Jackson Immunoresearch) in 1% donkey serum, 0.1% triton X-100, and 0.1% BSA in PBS for 3 hours at RT. After washing three times, cells were mounted onto coverslips using vectashield (Vectorlabs) before imaging. The randomly selected 10 areas of fluorescent images were taken, and the degeneration index was calculated by the ratio of the cumulative area of fragmented axons (defined as particles) to the intact axons using ImageJ.

**Amyloid beta treatment in mouse cortical neuron culture**

Embryonic cortices from E13.5-E14 ICR mice were dissected and collected in DPBS. To ensure the heterogeneity of neuronal population, cortices from at least 5 embryos were collected. Collected cortices were chopped with spring scissor and collected in 4.5 ml DPBS. The cortices were then incubated with 0.25% Trypsin by adding 500 µl of 2.5% trypsin solution. After 15 min incubation at 37 °C water bath, cells were triturated by mild pipetting. The reaction was stopped by adding 5 ml cell culture media, and the cells were collected by centrifugation at 440g for 3 min. The cell pellet was resuspended in 500 µl of cortical neuron culture media (Neurobasal with 2% B-27, 1% Glutamax, and 1% penicillin-streptomycin). The cells were stained with trypan blue and viable cells were automatedly counted by Countess II FL.

$2\times10^3$ cells were seeded to 8-well cell culture slide (SPL Life Science) coated with 100 µg/ml PDL. When DIV reached 3–5, cells were subjected to oligomeric Aβ treatment.

Oligomeric Aβ was prepared as previously described.[29] Briefly, recombinant Aβ (1–42) and reverse peptide (42–1) were purchased from Abcam and dissolved in DIW at 6 mg/ml concentration and stored at −80 °C. Peptide oligomerization was induced by incubating in DPBS at 1 mg/ml for 24 hours at 37 °C. Cortical neurons were then treated with 50 µg/ml of oligomeric Aβ for 24 hours in cortical neuron culture media. 1 µM recombinant mouse RBP4 or vehicle control was co-treated with Aβ.

The TUNEL assay was performed using ApopTag Red In situ Apoptosis Detection Kit (Merck Millipore) following the manufacturer's instructions. After TUNEL staining, the cells were incubated with 1:300 TuJ1 antibody in 1% donkey serum in PBS for 1 hour at RT. After washing thrice with PBS, 1:300 Alexa fluor 488-conjugated anti-rabbit antibody in 1% donkey serum in PBS was treated for 1 hour at RT. After Hoechst staining, the fluorescence image was taken from the randomly selected 7 areas, and at least 300 $TuJ1^+$ neuronal nuclei were counted for TUNEL co-staining.

**Determination of protein stoichiometry using single-molecule photobleaching assay**

DNA encoding DR6-Halo-His was generated by linking Halo-His tag to the C-terminus of DR6 ECD (1–349). DNA encoding RBP4-SNAP-His was generated by linking SNAP-His to the C-terminus of human RBP4 (1–201). DR6-Halo and RBP4-SNAP were inserted into the pLexm plasmid between EcoRI and XhoI restriction sites. For expression and purification, $3\times10^6$ SH-SY5Y cells were seeded in a 100 mm cell culture dish (SPL Life Science), and 7 µg plasmid was transfected using lipofectamine 2000. Cells were incubated for 4–5 days, and cultured media was collected and filtrated by a 0.45 µm cellulose acetate syringe filter. Filtrated media was directly incubated with pre-washed 300 µl of Ni-INDIGO agarose beads and 500 µM ATRA prepared in ethanol and incubated overnight at 4 °C with frequent agitation. The next day, Protein-bound resin was collected by centrifugation and briefly washed with 20 mM imidazole in PBS. Resins were then incubated with 1 ml of 5 µM SNAP-Surface Alexa Fluor 488 (New England Biolabs) in PBS with 1 mM DTT for SNAP labeling or 5 µM HaloTag ligand Alexa Fluor 660 (Promega) in PBS for Halo labeling for 1 hour at RT. After labeling, resins were collected by centrifugation and washed 4 times with 20 mM imidazole in PBS. Resin-bound proteins were then eluted in 50 µl of 500 mM imidazole in PBS (pH 7.4). Eluted DR6-Halo and RBP4-SNAP were mixed and incubated overnight at 4 °C. Labeled protein mixtures were used for photobleaching assay.

For stoichiometric determination of Alexa Fluor 488-labeled RBP4-SNAP and Alex Fluor 660-labeled DR6-Halo complex, the protein mixture was diluted to $1\times10^9$ particles/ml in PBS and then plated on 35 mm glass-bottom dishes (MatTek, P35G-0-10-C). The fluorescence intensities of individual DR6 and RBP4 molecules were determined by TIRF microscopy. The stoichiometry of the colocalized DR6-RBP4 complex was determined by the fluorescence intensities of each molecule.

**Single DR6 tracking on live cell membrane**

Highly inclined and laminated optical microscopy (HILO) was applied to image and track the DR6 at the apical plane of the cell using the TIRF system.

For quantum dot (QD) labeling of SNAP-DR6, U2OS cells stably expressing SNAP-DR6 or mutant were seeded in 35 mm glass-bottom dishes (MatTek) coated with collagen I (Santa Cruz Biotechnology). Cells were incubated with 1 nM benzylguanine-biotin (New England Biolabs) in cell culture media for 15 min at 37 °C and washed thrice with PBS. Next, Cells were incubated with 1 nM streptavidin-QD605 (Thermo Fischer Scientific) in cell culture media for 15 min at 37 °C and washed thrice with PBS. Cells were then incubated with 100 nM recombinant RBP4 or vehicle in cell culture media for 30 min at 37 °C. Before single-receptor tracking, media was removed and replaced with fresh cell culture media following a brief wash with PBS.

The video of QD signals were recorded for 1 min. The stacked images (20 ms time intervals) underwent two pre-processing steps: background subtraction and Gaussian filtering. To accurately determine the position of each particle at a finer scale than the pixel resolution, a two-dimensional cubic spline interpolation method was employed. The movement paths of the particles were determined by linking frame-to-frame positions using the Hungarian approach. For robust tracking, we set a minimum of 250 frames for track length and a maximum of 5 frames for closing gaps in the trajectory, ensuring longer and more precise tracks. The first ten points of the mean squared displacement (MSD) graph were used to compute MSD for each trajectory by following equation:

$$\mathrm{MSD}(\tau) = \langle r^2(\tau) \rangle = \langle |r_i(t+\tau) - r_i(t)|^2 \rangle = 4D\tau^{\alpha}$$

where $\tau$ is lag time, $r_i$ is a position of $i^{th}$ particle, D is diffusion coefficient, and $\alpha$ is an anomalous exponent. MATLAB v2020a was chosen for all these computational processes.

**HMM diffusive state analysis**

The transitions between two different states of diffusive movement in single DR6 molecules were examined using a Bayesian framework. This analysis was conducted with an unmodified HMM-Bayes algorithm as described in previous literature.[86] Custom MATLAB 2020a scripts facilitated all computational aspects of this analysis.

**Determination of DR6 oligomeric state using photobleaching assay**

$1\times10^5$ SH-SY5Y cells were seeded in a 24-well plate. The next day, 500 ng SNAP-DR6 plasmid was transiently transfected by lipofectamine 2000. After 48 hours, cells were treated with 5 µM SNAP-Surface Alexa Flour 488 in cell culture media for 30 min at 37 °C incubator. Cells were washed thrice by cell culture media and incubated with 1 µM recombinant RBP4 in PBS for 30 min at a 37 °C incubator. After incubation, cells were briefly washed twice with pre-chilled PBS and incubated with pre-chilled 4% PFA in PBS for 15 min on ice protected from light. Fixed cells were washed thrice by PBS, lysed in RIPA buffer, and homogenized by sonication. Cell lysates were diluted in PBS, and stoichiometric analysis was performed as described in DR6-RBP4 stoichiometric analysis.

**Calculation of IDR propensity and amino acid composition**

IDR was predicted by IUPred3 (https://iupred3.elte.hu)[87] and protein & RNA binding propensity was predicted by flDPnn (http://biomine.cs.vcu.edu/servers/flDPnn/).[56] The amino acid composition was profiled by a composition profiler (http://www.cprofiler.org).[88]

**Size exclusion chromatography of recombinant DR6 ECD**

Recombinant DR6 ECD was prepared in PBS without $CaCl_2$ at 1 mg/ml concentration. A Superdex 75 10/300 GL was filled with PBS with 1 mM TCEP. 17 or 50 µg recombinant DR6 ECD was diluted in 500 µl PBS, incubated for 1 hour at 4 °C, and directly injected for gel filtration. After collecting the whole fraction of DR6, the column was filled with PBS with 5 mM $CaCl_2$ and 1 mM TCEP. Recombinant DR6 ECD was diluted in 500 µl PBS with 5 mM $CaCl_2$, incubated for 1 hour at 4 °C, and directly injected for column running. 20 µl from each collected fraction was mixed with sample buffer with 10 mM DTT and boiled for 5 min at 95 °C. The equal volume of protein samples was loaded to 4–20% SDS-PAGE gel and analyzed by WB.

**Figures**

Biorender was used to make graphic schemes and ChimeraX[89] were used to make protein structure figures.

**Statistics and Reproducibility**

The statistical tests are performed by Graphpad Prism 9, as indicated in the figure legends. A minimum of 2 independent biological sample preparations or experiments were performed. Single-molecule analyses were performed by the randomized particle selection and unbiased data analysis. For experiments with purchased or purified recombinant proteins, the samples from at least two independent batches of protein purification were tested with consistent results.

**Data availability**

Proteomic data used to identify the proteins enriched by TurboID was deposited in **** and crosslinking mass spectrometry data was deposited in ****. Protein lists and crosslinked peptides are provided in Supplementary Table 1,2. The movie representing the tracking of QD-labeled DR6 is provided in Supplementary Movie 1–3. Full versions of all blots are provided in Supplementary Fig. 1. Further data and codes used in this paper can be received from the corresponding author on reasonable request. Source data are provided with this paper.

**Code availability**

Previously reported codes and prgrams were used to analyze photobleaching step analysis using HaMMy[90,91] and oligomerization dyamics anaylsis using HMM.[86,92,93].

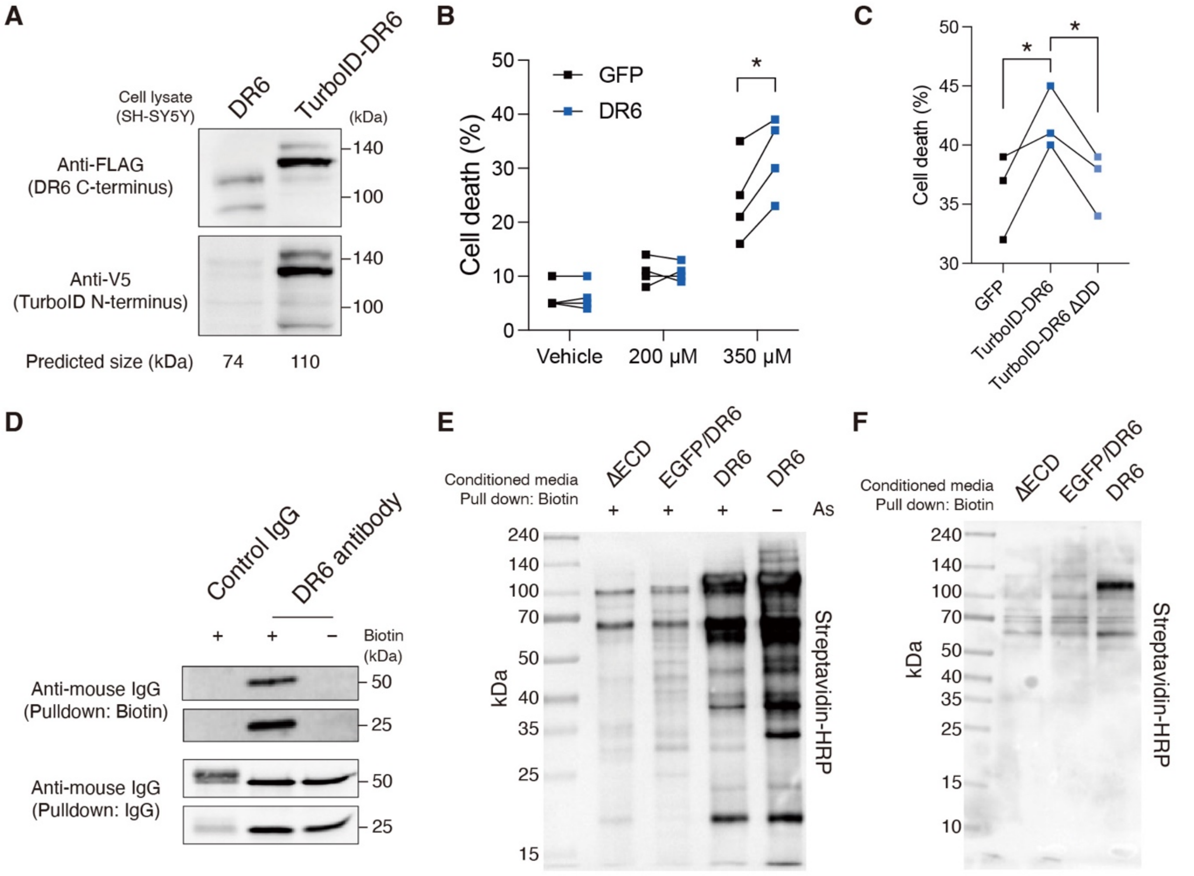


**Fig S1. Characterization of TurboID-DR6 and the protein biotinylation** (**A**) WB of DR6 or TurboID-DR6 expressed in SH-SY5Y detected by FLAG (DR6 C-terminus) or V5 (TurboID N-terminus) tag. (**B**) Cell death assay of SH-SY5Y expressing GFP or DR6 after 24 hours treatment of 0, 200, or 350 µM sodium arsenite (As). *$p$<0.05 by paired two-tailed t-test. (**C**) Cell death assay of SH-SY5Y expressing GFP, TurboID-DR6, or TurboID-DR6 Δdeath domain (ΔDD) after 24 hours treatment of 350 µM sodium arsenite. *$p$<0.05 by paired two-tailed t-test. (**D**) WB of biotinylated DR6 antibody collected by streptavidin beads after the induction of biotinylation. (**E**) WB of the proteins biotinylated during 350 µM sodium arsenite treatment for 24 hours. (**F**) WB of biotinylated proteins collected from the cultured media of Hs683 expressing TurboID-DR6 or controls.

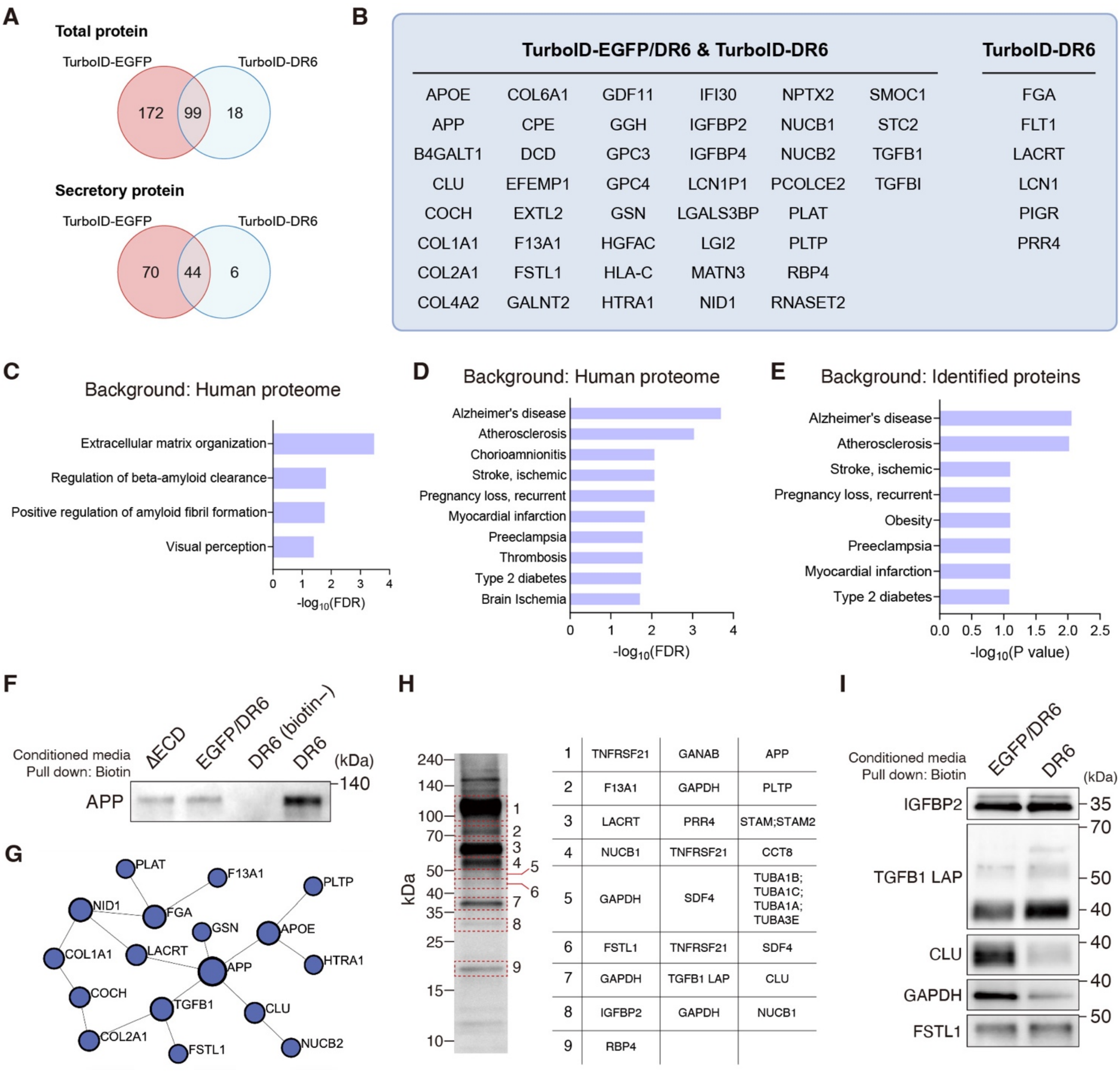


**Fig. S2. See next page for caption**

**Fig S2. DR6-associated secretome projects the proteomic landscape of AD** (**A**) Venn diagram showing the number of total or secretory proteins identified by LC-$MS^2$ in the cultured media from SH-SY5Y expressing either TurboID-EGFP/DR6 or TurboID-DR6. (**B**) The list of secretory proteins found in both TurboID-EGFP/DR6 and TurboID-DR6 or only in TurboID-DR6 sample. (**C** to **E**) Gene ontology (GO) analysis (C) and disease association over human proteome (D) or identified secretory proteins (E) of 50 proteins listed in (B). False discovery rate (FDR) was used to adjust the multiple comparisons (**F**) WB of APP in biotinylated proteins. (**G**) Network analysis of the secretory proteins enriched by TurboID-DR6. (**H**) The list of top-abundant proteins in each gel cut. (**I**) WB of AD-associated proteins in the biotinylated proteins by TurboID-EGFP/DR6 or TurboID-DR6.

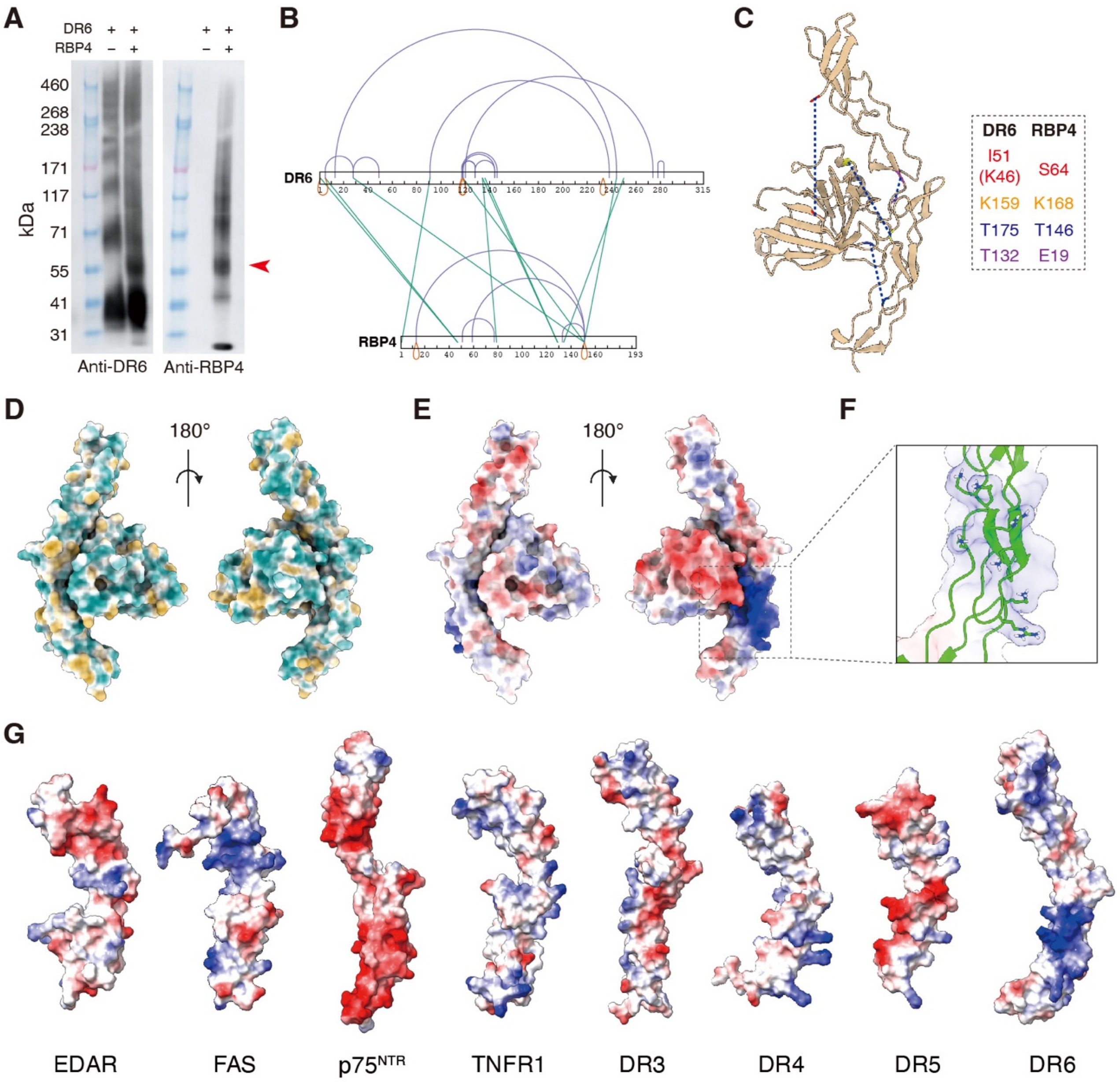


**Fig S3. Crosslinking mass spectrometry and docking simulation of RBP4-DR6 complex** (**A**) WB of DR6 (left) and RBP4 (right) after crosslinking by $BS^3$. Red arrow indicates the 1:1 crosslinked complex which was subjected to LC-$MS^2$. (**B**) Mapping of crosslinked sites. (**C**) Graphic view of crosslinked sites used for protein docking simulation. (**D** and **E**) Graphic over-view of hydrophobic (D) and charged (E) surface of DR6-RBP4 complex. (**F**) Close view of the DR6 positively charge surface contacting RBP4. (**G**) Graphic overview of charged surface of cysteine-rich domains (CRDs) of DRs predicted by Alphafold.

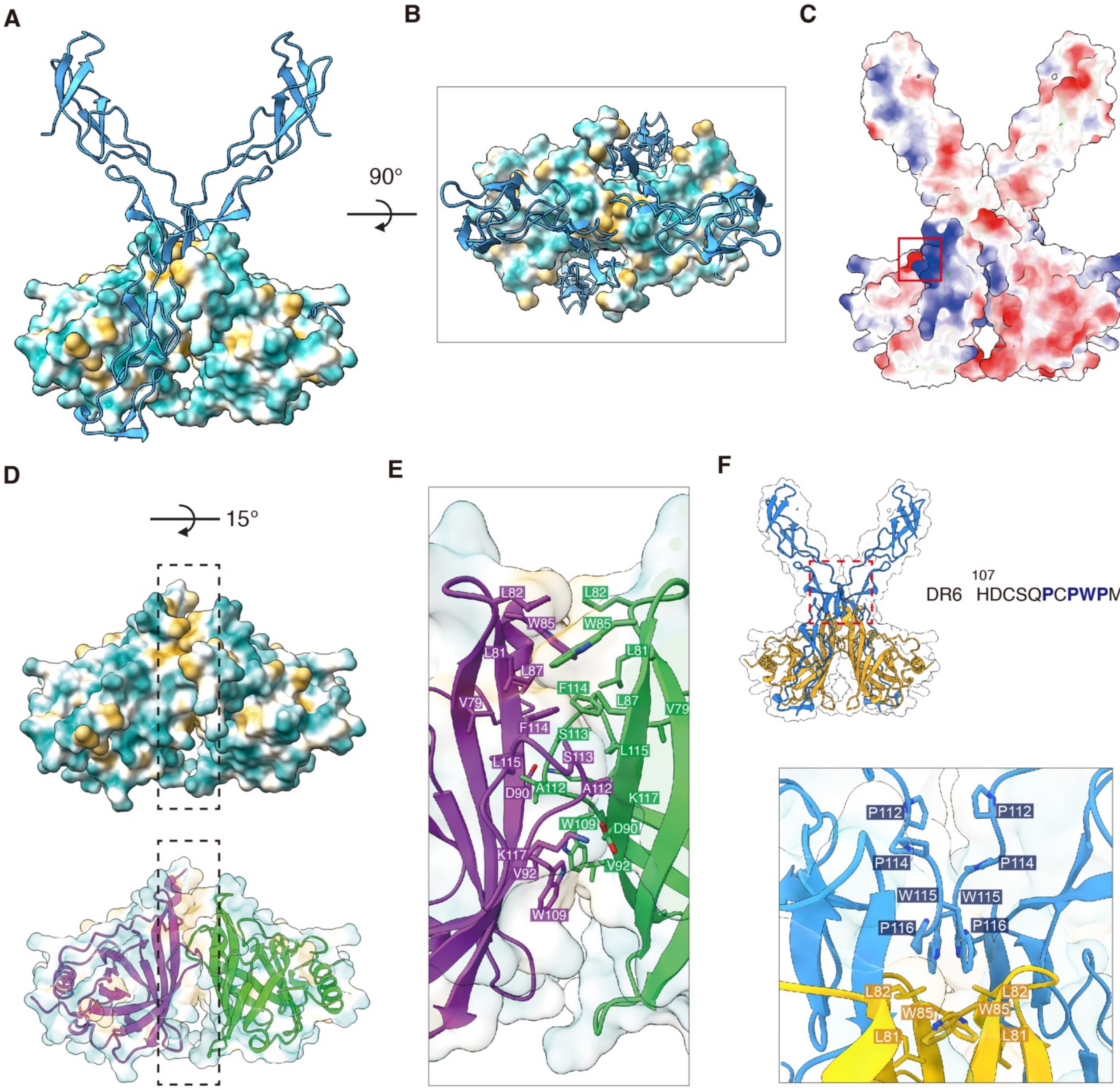


**Fig S4. Structural property of RBP4-DR6 complex predicted by Alphafold3.** (**A**) Graphic showing hydrophobic RBP4-RBP4 interaction in complexed with DR6 (green). Surface color indicates the hydrophobicity of RBP4. (**B**) Top-view of RBP4-DR6 complex. (**C**) Polar surface of RBP4-DR6 complex. DR6 positively charged surface participates in the interaction with RBP4. Red rectangle indicates the binding site depicted in Figure 2H. (**D**) RBP4-RBP4 hydrophobic binding interface. (**E**) Close view of RBP4-RBP4 binding interface. Hydrophobic residues are symmetrically aligned. (**F**) Additional alignment of DR6 hydrophobic residues within the RBP4-DR6 complex.

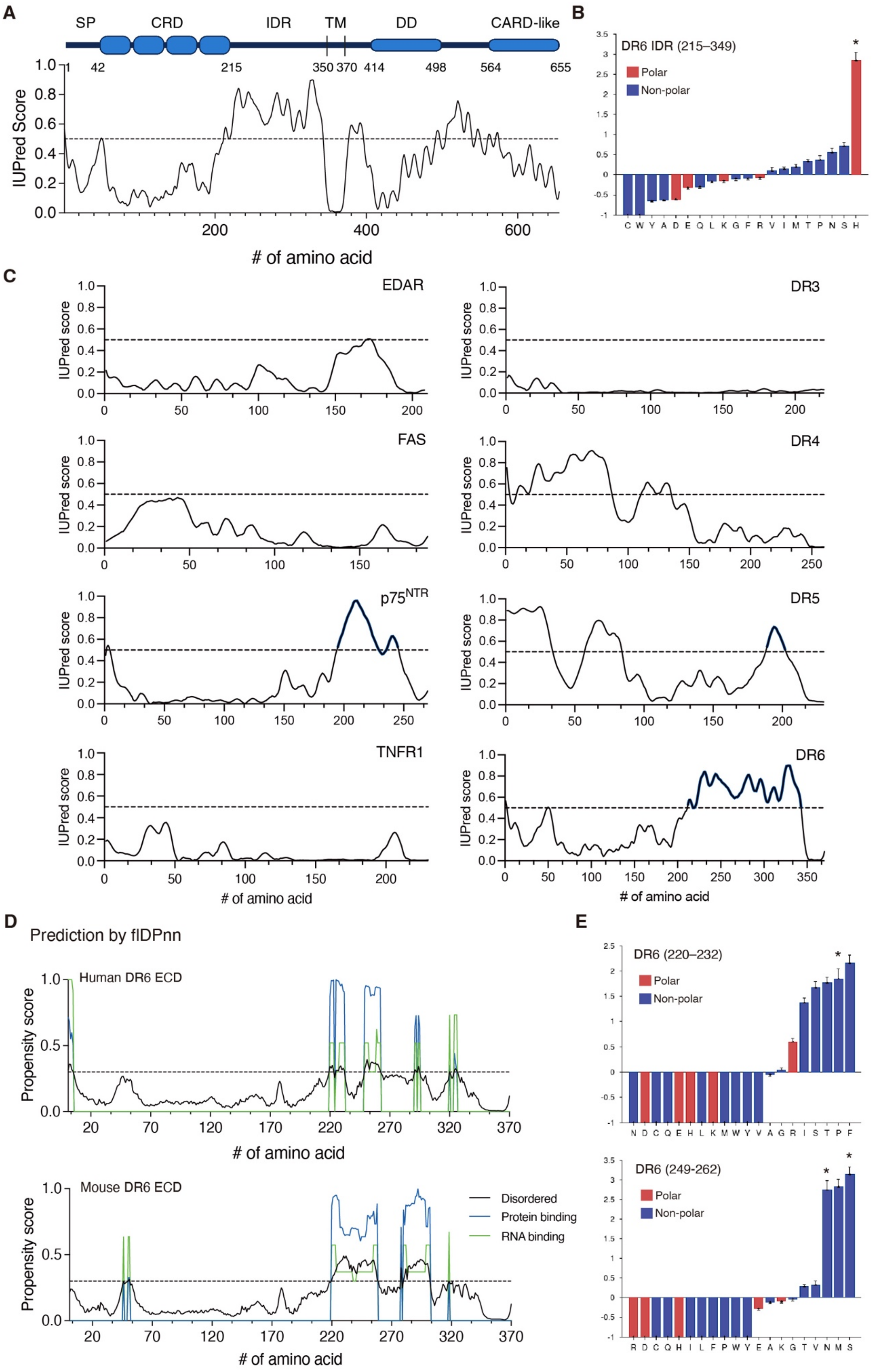


**Fig. S5. See next page for caption**

**Fig S5. Information on the DR6 IDR and its amino acid composition** (**A**) IUPred score of full-length human DR6 and a graphic showing domain architecture. (SP, signal peptide; CRD, ysteine-rich domain; IDR, intrinsically disordered region; TM, transmembrane domain; DD, death domain; CARD-like, caspase recruitment domain-like domain) (**B**) Amino acid composition profile of DR6 IDR (215–349) calculated by composition profiler over Disprot 3.4.[94] (**C**) Comparison of IUPred score of DR ECDs. Blue line indicates the regions predicted to be disordered (>0.5). (**D**) Propensity score for disordered, protein-binding, or RNA-binding propensity of mouse and human DR6 ECD calculated via flDPnn. The dotted line indicates the threshold for the disordered region (>0.3). (**E**) Composition profile of DR6 short IDRs (220–232 and 249–262) calculated by composition profiler over Disprot 3.4. *P<0.05 was evaluated using relative entropy as previously described.[88]

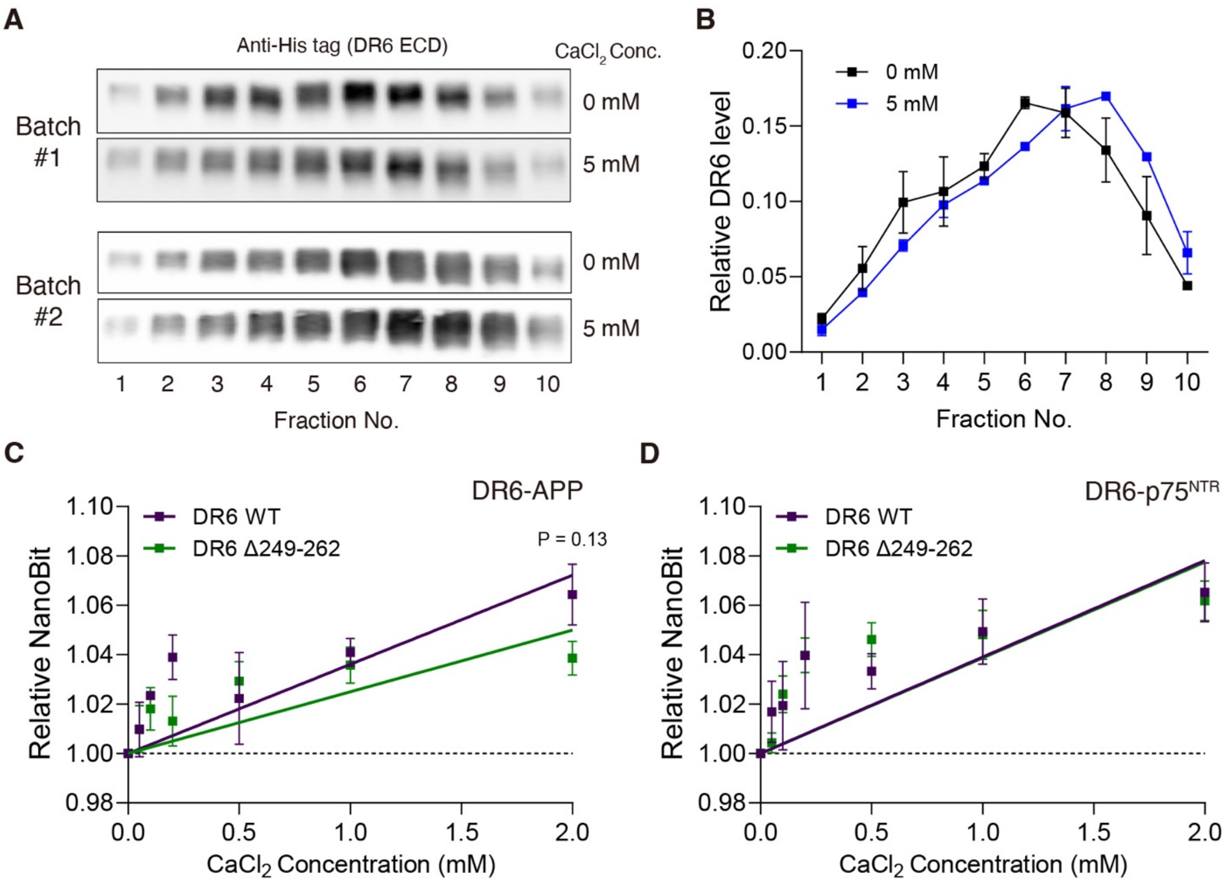


**Fig S6. Effect of Calcium on DR6 oligomerization and heterotypic interaction** (**A**) WB (anti-his tag) of recombinant DR6 ECD-His separated by size exclusion chromatography with or without $CaCl_2$ (5 mM) in running buffer (PBS). (**B**) Quantification of relative band intensity of DR6 ECD separated by size exclusion chromatography (n=2). Data are presented as mean ± SD determined from independent protein batches and experiments. (**C** and **D**) NanoBit assay measuring the effect of DR6 IDR (249–262) on the elevating concentration of $CaCl_2$ on the interaction of DR6 with APP (C; n=5) or $p75^{NTR}$ (D; n=4). DR6 WT data presented in (Figure 5H) was compared with DR6 Δ249–262. Data are presented as mean ± SEM determined from independent experiments performed in quadruplicate. The P-value was calculated by unpaired two-sided t-test.

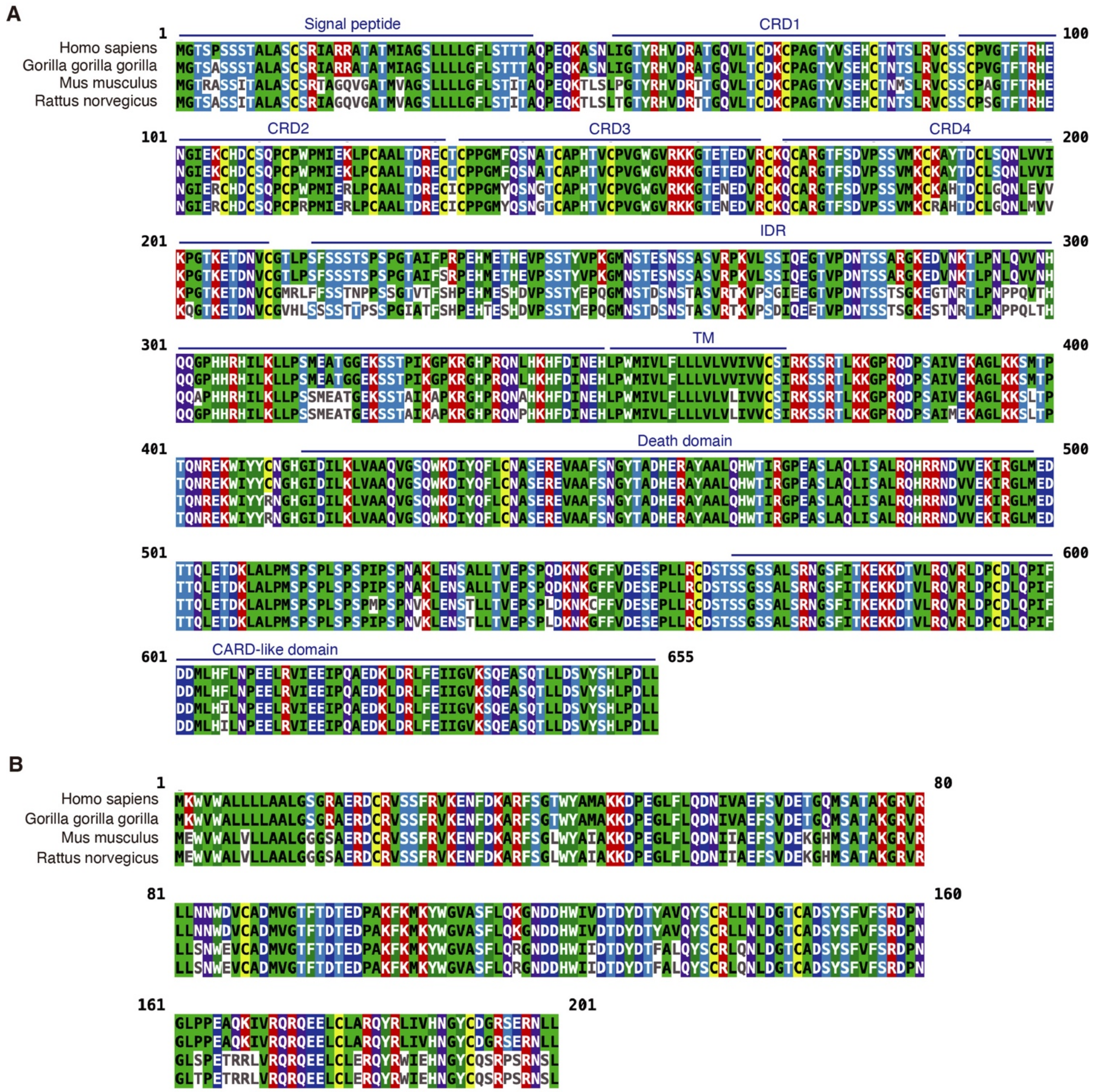


**Fig S7. Sequence conservation and homology of DR6 and RBP4 within Vertebrata.** (**A**) Comparison of DR6 sequence between human and model organisms. Structural domains are indicated above the sequence. (**B**) Comparison of RBP4 sequence between human and model organisms.

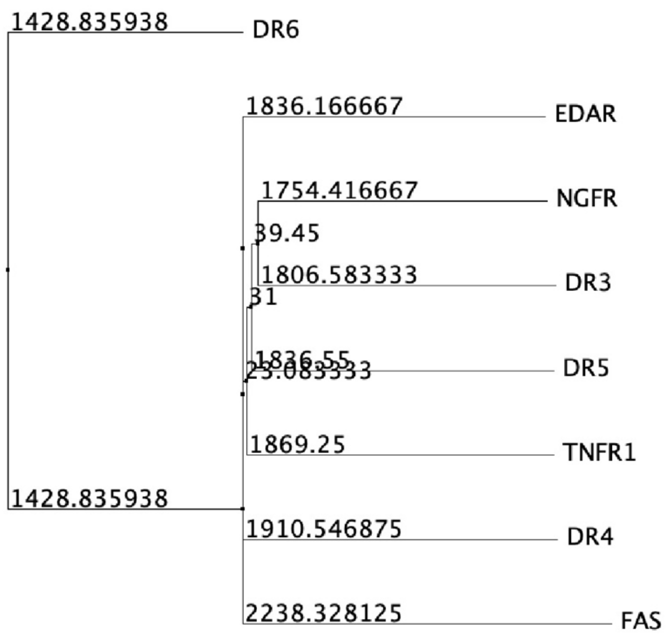


**Fig S8. DR6 is distant from other DRs.** The average distance tree of human DRs using BLOSUM62 shows the structural distance of DR6. The number indicates the distance.
